# Global Protein-Turnover Quantification in *Escherichia coli* Reveals Cytoplasmic Recycling under Nitrogen Limitation

**DOI:** 10.1101/2022.08.01.502339

**Authors:** Meera Gupta, Alex Johnson, Edward Cruz, Eli Costa, Randi Guest, Sophia Hsin-Jung Li, Elizabeth M Hart, Thao Nguyen, Michael Stadlmeier, Benjamin P. Bratton, Thomas J. Silhavy, Ned S. Wingreen, Zemer Gitai, Martin Wühr

## Abstract

Protein turnover is critical for proteostasis, but turnover quantification is challenging, and even in well-studied *E. coli*, proteome-wide measurements remain scarce. Here, we quantify the degradation rates of ∼3.2k *E. coli* proteins under 12 conditions by combining heavy isotope labeling with complement reporter ion quantification and find that cytoplasmic proteins are recycled when nitrogen is limited. We use knockout experiments to assign substrates to the known cytoplasmic ATP-dependent proteases. Surprisingly, none of these proteases are responsible for the observed cytoplasmic protein degradation in nitrogen limitation, suggesting that a major proteolysis pathway in *E. coli* remains to be discovered. Lastly, we show that protein degradation rates are generally independent of cell division rates. Thus, we introduce broadly applicable technology for protein turnover measurements and provide a rich resource for protein half-lives and protease substrates in *E. coli*, complementary to genomics data, that will allow researchers to decipher the control of proteostasis.

## Introduction

Protein degradation is central to protein homeostasis (proteostasis) and is critical in most cellular pathways.^1-3^ As environments change, modification of degradation rates can rapidly adapt protein abundances to desired levels. Even if protein levels are modulated via transcription or translation, a protein’s response time is set by its replacement rate.^4,5^ Unsurprisingly, many signaling and transcriptional regulatory proteins exhibit short half-lives.^6-9^ Protein degradation is important in health and disease, such as cancer and neurodegenerative disorders.^10,11^ Additionally, protein degradation plays an important metabolic role. It has been shown that bacteria and yeast cells increase their proteome turnover rates under starvation conditions, presumably generating and recycling scarce amino acids.^12-15^

Quantitative models have been developed to describe the dependence of global protein expression on cells’ physiological characteristics, most notably cell doubling times. These models are probably the most well-developed for the model bacterium *Escherichia coli*.^16,17^ The cell cycle time in *E. coli* varies from 20 minutes in rich media to the cessation of division under starvation. Transcription rates typically increase with cell division rates.^18,19^ Knowing how global parameters scale with physiological cell states allows for remarkable quantitative predictions for gene expression changes across different growth conditions.^16,17^ However, active protein degradation by proteolysis is typically ignored in these models. Instead, proteins are assumed to be completely stable and only diluted via cell growth and division. This simplification is likely due to a lack of reliable genome-wide degradation rate measurements under varying growth conditions. It is still unclear how active degradation rates scale with changing cell cycle times and how this affects global gene expression regulation. Knowledge of protein degradation rates and how they scale with the physiological characteristics of cells would improve predictive models of protein expression across various cell states.

Cells have developed sophisticated mechanisms to recognize and degrade specific proteins. While eukaryotes utilize the ubiquitin-proteasome pathway, in bacteria, selective proteolysis is executed by ATP-dependent proteases.^1^ While many proteases can digest unfolded proteins and peptides, unfolding a protein for degradation requires energy. In *E. coli*, four ATP-dependent proteases are known: ClpP, Lon, HslV, and FtsH. Pulldown experiments with inactivated protease mutants or protein-array studies have allowed the proteome-wide identification of putative substrates.^20-23^ Orthogonally, individual substrates have been assigned to the four proteases by measuring the degradation of individual proteins in protease knockout strains or via *in vitro* assays.^24,25^ Several example proteins (e.g., RpoH, LpxC, and SoxS) have been shown to be degraded by multiple proteases, demonstrating remarkable redundancy.^26-28^ But it is still unclear to what extent substrates overlap between different proteases.

In most biological systems, protein degradation is balanced by the synthesis of new protein, making measurements of degradation rates challenging. An easy way to overcome this complication is by using translational inhibitors like cycloheximide or chloramphenicol.^29-31^ Assuming that the addition of the drug does not perturb the cells aside from blocking the translation of new proteins, protein degradation can be conveniently measured by assaying changes in protein abundances over time via western blots or quantitative proteomics. However, when we performed such experiments in *E. coli*, we found that many proteins whose abundances rapidly decreased were periplasmic (Fig. S1). Further investigation revealed that these periplasmic proteins were not degraded but rather were accumulating in the bacterial growth medium (Fig. S1). Presumably, this was due to protein leakage through the outer membrane. We concluded that translation inhibitor experiments in *E. coli* could lead to major perturbations, and, thus, interpreting such studies might be challenging.

A classic method to measure the unperturbed turnover of biological molecules uses radioactive isotope tracking or the combination of heavy isotope labeling and quantitative mass spectrometry.^32,33^ Isotopic labels can be introduced with heavy nutrients (e.g., ammonia, glucose, or amino acids) or by incubation in heavy water. Most proteomic turnover studies have been performed with heavy amino acid labeling (dynamic SILAC)^34,35^, but the small number of labeled residues limits sensitivity for short-time SILAC labeling, and missing values can hinder the coverage of multiple time points in complex systems. A further advance has been the combination of SILAC experiments and isobaric tag labeling.^36^ However, these measurements tend to suffer from the inherent ratio compression of multiplexed proteomics.^37-39^ Heavy ammonia, glucose, and water are comparatively cheap but result in overly complex MS1 spectra, which are difficult to interpret, particularly for lower abundance proteins.^40-42^ For a more detailed discussion of the advantages and limitations of various global protein turnover measurement techniques, please see the recent review by Ross et al., particularly Table S1.^43^

Despite the central role of protein degradation in nearly every aspect of biological regulation, reliable and large-scale measurements are still scarce. Even fewer studies have compared degradation rates between multiple conditions.^44,45^ Here, we measure protein turnover in *E. coli* by combining heavy isotope labeling via ^15^N ammonia with the accurate multiplexed proteomics method TMTproC.^46^ We provide a rich resource of protein degradation rates for ∼3.2k *E. coli* proteins (77% of all genes in *E. coli*) measured across 12 different growth conditions with replicates. When comparing degradation rates among various nutrient limitations, we found that *E. coli* recycles its cytoplasmic proteins when nitrogen-limited, and we assign substrates to proteases by measuring the change of protein turnover in knockout strains. Lastly, we show that active degradation rates are typically independent of cell division rates.

## Results

### Combining heavy isotope labeling with complement reporter ion quantification enables high-quality protein turnover measurements

We wanted to measure protein degradation rates and evaluate how these rates vary across growth conditions. To simplify our measurements, we grow *E. coli* in chemostats, where we can control the cell doubling time and enforce steady state (Fig. 1A). After cells reach steady state, we change the inlet medium from unlabeled nutrients to ^15^N-labeled ammonia. Over time, the ^15^N-ammonia concentration in the reactor increases, and newly synthesized proteins incorporate more heavy isotopes. We can monitor the shift in the isotopic envelope of peptides by taking samples after the media switch using proteomics (Fig. 1B). With the knowledge of a peptide’s chemical composition and the fraction of heavy isotopes over time, we can calculate the degradation rate of the corresponding protein. In practice, however, obtaining such measurements of isotopic envelopes in the MS1 spectrum is quite challenging, particularly at later time points when the isotopic envelopes spread out and overlap with those of other peptides. Additionally, missing values between time points are a severe limitation of such approaches.^42^ To overcome these limitations, we labeled samples at each of the acquired eight time points with TMTpro isobaric tags and combined them for co-injection into the mass spectrometer.^46,47^ Analyzing these extremely complex samples with standard low *m*/*z* reporter ion quantification would lead to severe ratio distortion and measurement artifacts.^37,38,48^ We overcame this limitation by isolating the pseudo-monoisotopic peak (M0) for fragmentation and quantifying the complement reporter ions in the MS2 (Fig. 1C).

**Figure 1:**
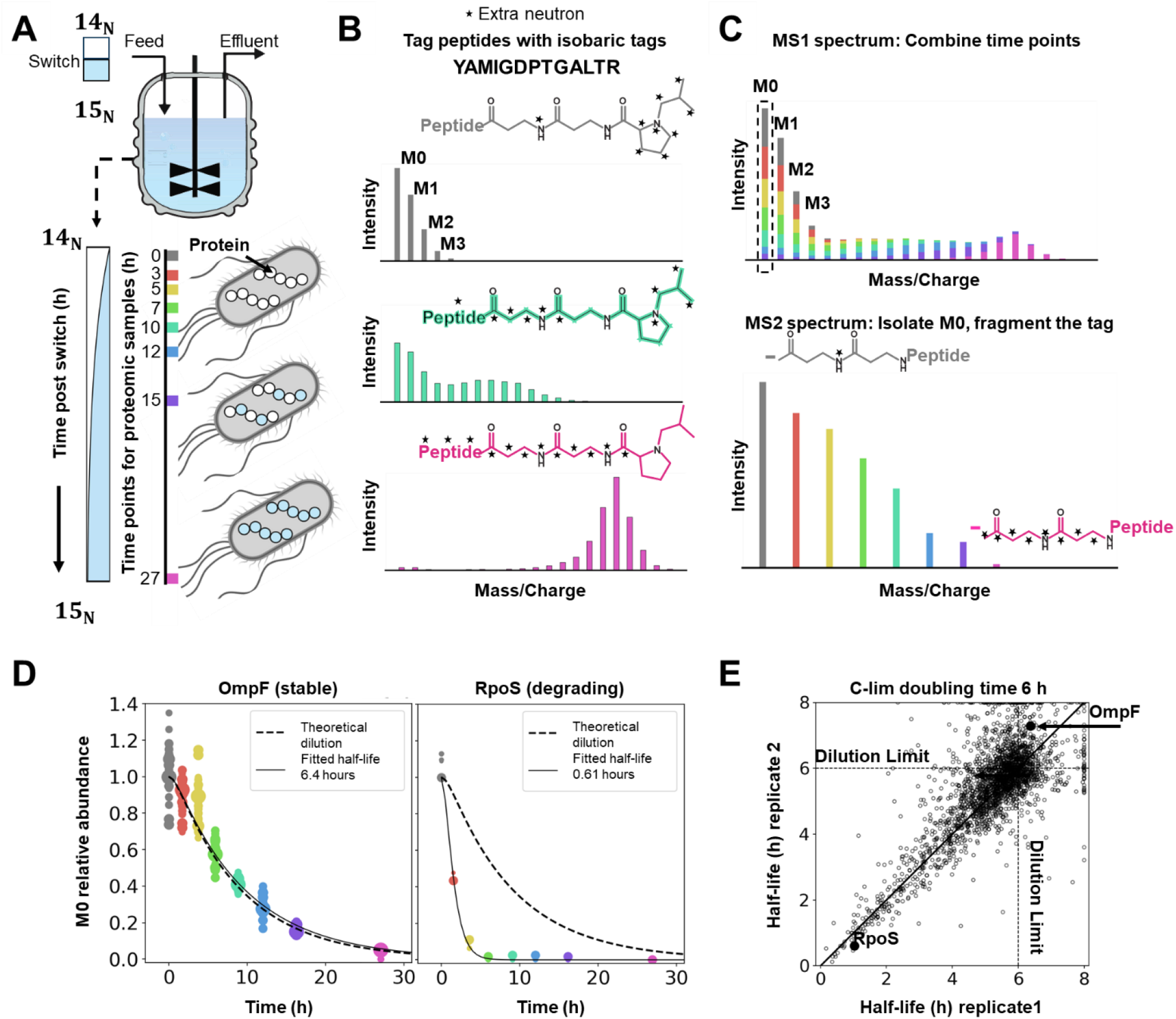
Combining heavy isotope labeling with an accurate multiplexed proteomics method (TMTproC) enables high-quality measurements of unperturbed protein turnover. **A)** Experimental setup. *E. coli* cells were grown in chemostats with a defined doubling time. After reaching steady state, the chemostat feed was switched to a medium with _15_N-labeled ammonia. Newly synthesized proteins will increasingly incorporate heavy isotopes. Proteomics samples were collected at various time points to determine the protein turnover rate. **B)** Theoretical isotopic envelopes of an example tryptic peptide, which is assumed to be stable (protein is removed from the vessel only through dilution). Over time, the increasing fraction of heavy ammonia in the peptide’s structure shifts the isotopic envelope to higher masses. Peptides were labeled with isobaric tags (TMTpro) to encode different time points. **C)** Top: theoretical MS1 spectrum for a single peptide species after combining labeled peptides from all the time points. The mass spectrometer was set to isolate the monoisotopic peak (M0) and fragment the peptide. Bottom: the resulting complement reporter ions (peptide plus broken tag) enable accurate quantification of the relative abundance within the M0 peak over time. **D)** Example measurements for the stable OmpF protein and rapidly degrading RpoS protein. Each dot indicates the peptide quantification relative to the median level measured when the feed was switched. The size of each point is proportional to the number of measured ions. Fitting the observed data with the theoretical decay profile for M0, we can extract the half-life for each protein (solid curve). The dotted curve shows the theoretical decay for a stable protein. **E)** Scatter plot of measured protein half-lives for biological replicates of carbon-limited *E. coli* grown with a 6-hour doubling time. Dotted lines indicate the cell doubling times. The solid line marks the 1:1 line. The half-lives for each protein were calculated from the fits shown in D. Median standard deviation for the half-lives between the replicates is 0.3h.

Figure 1D shows the peptide-level quantification for a stable protein, OmpF, and an unstable protein, RpoS, from carbon-limited chemostats with 6-hour doubling times.^24,49,50^ Proteins without active degradation are expected to follow the theoretical dilution curve (dotted curve) based on the chemostat dilution rate. Fitting the measured signal of OmpF peptides with a model for the expected decay of the M0 peak (solid curve) results in a turnover half-life similar to this expected value. In contrast, the deduced half-life for RpoS is much shorter than the cell doubling time. We obtain half-lives for ∼2.6k *E. coli* proteins per experiment with a median standard deviation of 0.3 hours (Fig. 1E, Table S1). Having established this technology, we acquired similar measurements for 12 different growth conditions, each with two biological replicates, quantifying the degradation rates of ∼3.2k proteins in at least one condition (Table 1, Table S1). We then used this resource to investigate how *E. coli* adapts protein turnover under various growth conditions.

**Table 1:** Summary of the 12 growth conditions for which we measured protein turnover rates.

|  | Strain<br>NCM3722 | Condition | Reactor<br>type | Doubling<br>Time | Replicates | # of<br>proteins |
| --- | --- | --- | --- | --- | --- | --- |
| 1. | Wild Type | Minimal<br>media | Batch | 42 mins | 2 | 2555 |
| 2. | Wild Type | C-lim | Chemostat | 3 h | 2 | 2665 |
| 3. | Wild Type | C-lim | Chemostat | 6 h | 2 | 2651 |
| 4. | Wild Type | C-lim | Chemostat | 12 h | 2 | 2697 |
| 5. | Wild Type | P-lim | Chemostat | 6 h | 2 | 2469 |
| 6. | Wild Type | P-lim | Chemostat | 12 h | 2 | 2619 |
| 7. | Wild Type | N-lim | Chemostat | 6 h | 2 | 2467 |
| 8. | Wild Type | N-lim | Chemostat | 12 h | 2 | 2460 |
| 9. | $\Delta$ hslV | N-lim | Chemostat | 6 h | 2 | 2393 |
| 10. | $\Delta$ lon | N-lim | Chemostat | 6 h | 2 | 2390 |
| 11. | $\Delta$ clpP | N-lim | Chemostat | 6 h | 2 | 2491 |
| 12. | $\Delta$ hslV $\Delta$ lon<br>$\Delta$ clpP | N-lim | Chemostat | 6 h | 2 | 2809 |
|  |  |  |  |  | **Union** | **3252** |

### *E. coli* recycles its cytoplasmic proteins under nitrogen limitation

Building on our method to measure protein turnover, we wanted to compare protein degradation rates under various nutrient limitations. To this end, we compared carbon (C-lim), phosphorus (P-lim), and nitrogen (N-lim) limitation measurements from chemostats with 6-hour doubling times. We found that most proteins in C-lim are stable with a measured total half-life close to the theoretical dilution time (Fig. 2A). Using biological replicates to identify degrading proteins with high confidence (Fig. 2B), we found that 15% of the proteome is actively degraded in C-lim (*p*-values < 0.05). Protein half-lives under P-lim have a similar distribution and a similar percentage of proteins that degrade with high confidence. However, in N-lim we found that 43% of proteins are actively degraded (Fig. 2B, *p*-value < 0.05).

**Figure 2:**
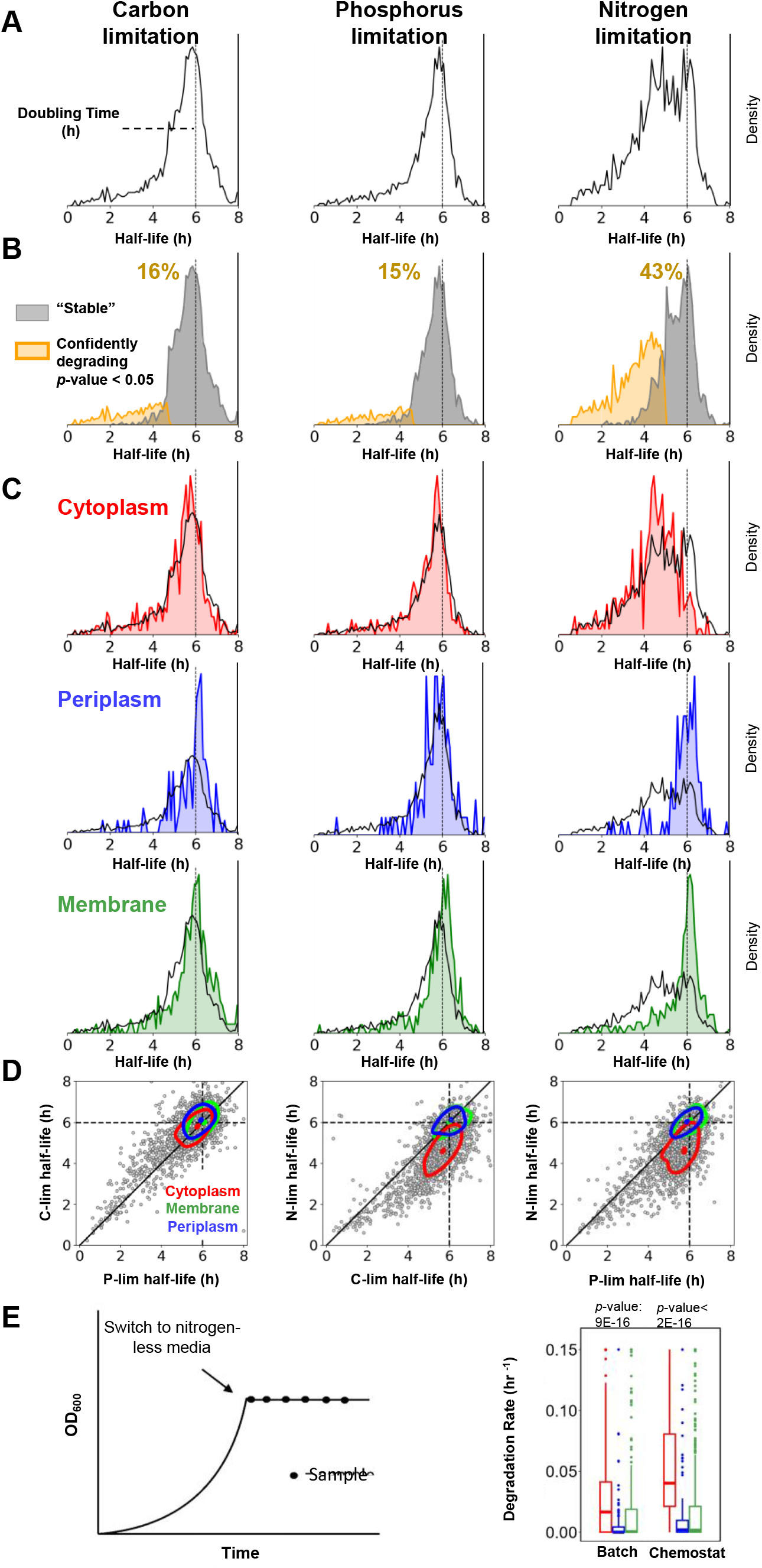
*E. coli* recycles its cytoplasmic proteins when nitrogen is limited. **A)** Histogram of protein half-lives for *E. coli* grown in chemostats under C-lim, P-lim, and N-lim. The vertical line marks the dilution limit set by the 6h doubling time. Half-lives greater than doubling time indicate measurement noise. Under C-lim and P-lim, most proteins have half-lives equal to the doubling time, suggesting they are stable. However, under N-lim many proteins are actively degraded. **B)** Separation of the proteome into stable proteins (grey) and actively degrading proteins (yellow) (*t*-test, *p*-value < 0.05, *n*=2). In C-lim and P-lim, 15% of the proteome turns over with high confidence. In contrast, under N-lim, 43% of the proteome turns over. **C)** Distribution of half-lives for proteins from different subcellular localizations overlayed against the entire proteome. Most proteins are stable under C-lim and P-lim, irrespective of localization. However, nearly all cytoplasmic proteins slowly degrade under N-lim while the membrane and periplasmic proteomes are largely stable. **D)** Scatter plots of protein half-lives in different nutrient limitations. The dotted black lines mark the dilution limit, the solid black line denotes perfect agreement. Contour plots contain 50% of the probability mass for each subcellular compartment. The contour plots of membrane and periplasmic proteins are centered around the dilution limit in all the binary comparisons, indicating that most of these proteins are stable under all limitations. However, the shift in the contour plots of the cytoplasmic proteins on comparing N-lim with P-lim and C-lim suggests that the cytoplasmic proteins are degraded in N-lim. **E)** Measurement of protein decay rates under complete nutrient starvation in batch. Exponentially growing cells in minimal media are washed and resuspended in nitrogen-depleted media. Proteomics samples are collected after the switch and protein profiles are fitted with exponential curves to obtain the decay rates. In batch, similar to the N-lim chemostat, the cytoplasmic proteins are decaying with high confidence as compared to the membrane and periplasmic proteins (ANOVA).

We found that the increase in protein degradation in N-lim could be attributed to the active degradation of a wide range of cytoplasmic proteins. The mode protein half-life for membrane and periplasmic proteins in all three conditions is very close to the theoretical dilution limit. In contrast, the mode protein half-life for cytoplasmic proteins is significantly shorter under N-lim than C-lim or P-lim. We estimate that 56% of cytoplasmic proteins are actively degraded in N-lim, while only 13% of membrane proteins and 4% of periplasmic proteins undergo active degradation in this condition. Due to measurement noise, these estimates are likely lower bounds of the true extent of protein degradation in N-lim.

We then tested whether cytoplasmic protein degradation in N-lim chemostats extends to the more physiologically relevant case of batch starvation. We grew *E. coli* cells in minimal medium until they reached an OD of ∼0.4. We then switched the exponentially growing cells into medium depleted of nitrogen (Fig. 2E). Once again, many cytoplasmic proteins are degraded under nitrogen starvation, and membrane/periplasmic proteins are largely stable. Thus, *E. coli* cells slowly degrade their cytoplasmic proteins when nitrogen is scarce in both chemostats and batch cultures. About 2/3 of the cell’s nitrogen is stored in proteins.^51^ The degradation of proteins upon nitrogen starvation likely allows the regeneration and recycling of scarce amino acids and enables *E. coli* to produce new proteins to adapt to new environments.

### Measuring protein turnover in knockout mutants allows the identification of protease substrates

Next, we were interested in discovering the protease(s) responsible for the large-scale turnover of cytoplasmic proteins in N-lim. Combining protein-turnover measurements with genetic protease knockouts allows us to investigate protease-substrate relationships on a proteome-wide level. Since unfolding and degrading stably folded cytoplasmic proteins requires energy, we focused on assigning substrates to the ATP-dependent proteases. In *E. coli*, there are four known cytoplasmic ATP-dependent protease complexes: ClpP (in complex with ClpX or ClpA), Lon, HslV (in complex with HslU), and FtsH.^1^ We identify putative substrates for the first three of these proteases by comparing the protein half-lives in protease knockout (KO) with wildtype (WT) cells (Fig. 3A). We were able to validate several known protease-substrate targets and identify novel degradation pathways using these experiments. For example, the unfoldase ClpA is completely stabilized by knocking out *clpP*, consistent with previous literature.^52^ We identified Tag and UhpA as putative novel substrates of Lon and HslV, respectively. However, many proteins still degrade in the three protease KO lines, e.g., the phosphatase YbhA—which contributes to Vitamin B6 homeostasis—still rapidly turns over with a half-life of ∼1 hour in each knockout strain.^53^ Surprisingly, even the proteins that increase their half-lives in single KOs are often not completely stabilized. Additionally, bulk cytoplasmic proteins are still degraded in all three single KOs.

**Figure 3:**
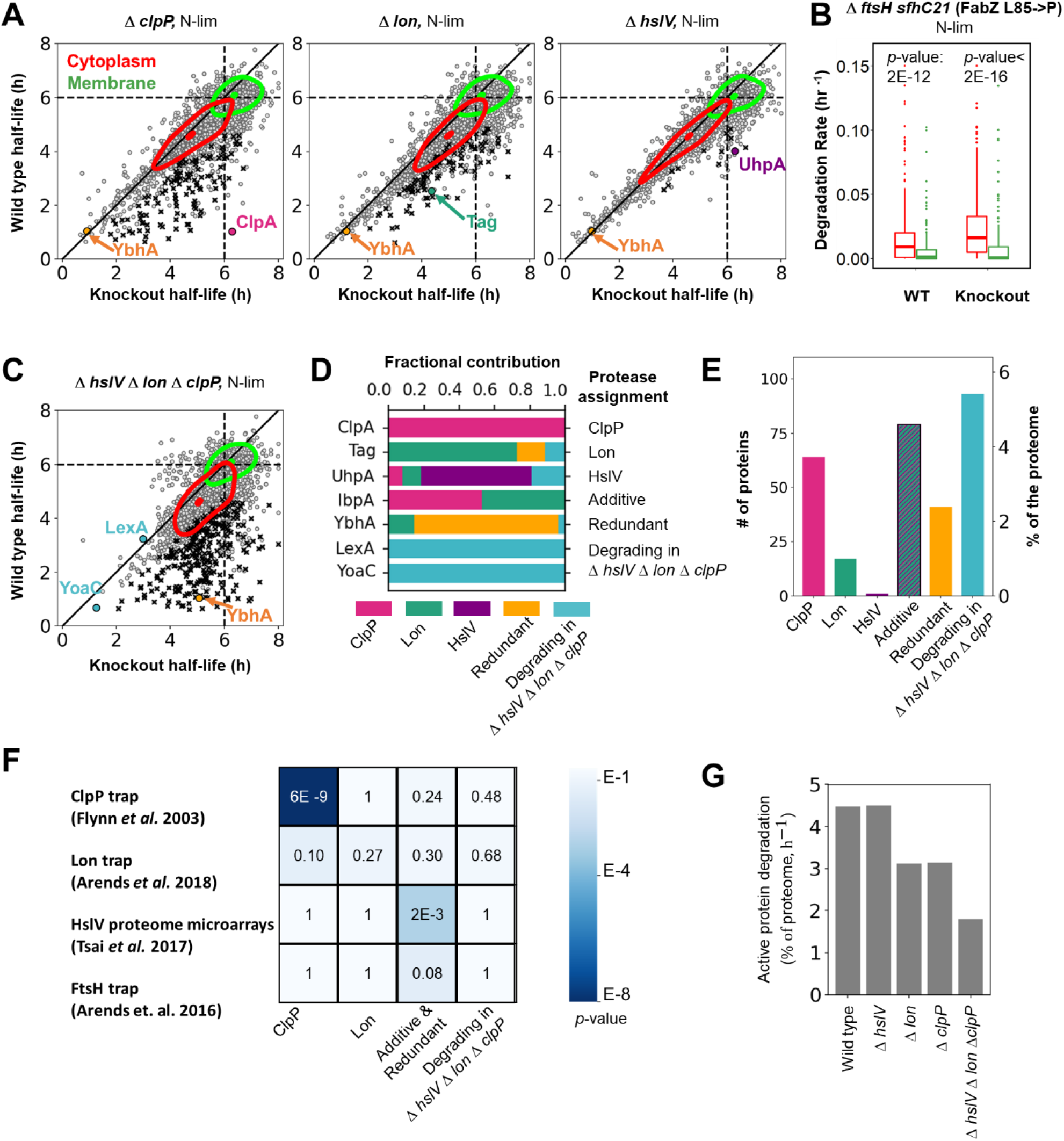
Measurement of protein turnover in protease knockout strains enables proteome-wide identification of protease substrates. **A)** Scatter plots of protein half-lives of N-limite il type (WT) compare to Δ*clpP*, Δ*lon*, and Δ*hslV* knockout (KO) cells. Dotted lines mark the dilution limit, and the solid black line indicates perfect agreement. Substrates, marked in black, increase their half-lives in KOs with high confidence (*t*-test, *p*-value < 0.10). ClpA (in pink), Tag (in teal), and UhpA (in purple) are the substrates of Δ*clpP*, Δ*lon*, an Δ*hslV*, respectively. However, the protein YbhA (in orange) is still degraded in the individual KOs. Contour plots containing 50% of the probability mass for the cytoplasmic (in red) and membrane (in green) proteins indicate that individual KOs of *hslV, lon*, and *clpP* retain their ability to degrade the bulk cytoplasmic proteins. **B)** ince Δ*ftsH* cells cannot grow in chemostats, we repeated the batch starvation assay as in Fig. 3E. Cytoplasmic proteins are shown in red and membrane proteins in green. Results in icate that the Δ*ftsH* cells, like the WT, also degrade their cytoplasmic proteins under nitrogen starvation (*t*-test, *p*-value = 2E-12 for WT and *p*-value < 2E-for Δ*ftsH*). **C)** Scatter plots of protein half-lives of WT and Δ*clpP* Δ*lon* Δ*hslV* cells in N-lim. The substrates marked in black increase their half-lives in the triple KO (*t*-test, *p*-value < 0.10). Strikingly, many proteins are still degrading in the triple KO, e.g., LexA and YoaC in blue. In fact, the bulk cytoplasm is still turning over. However, many more proteins are stabilized in the triple KO compared to the individual KOs, indicating redundancy among substrates. E.g., YbhA, which is still degrading in the individual knockouts, gets significantly stabilized in the triple KO (in orange). **D)** Comparing the shifts in the WT and KO strains’ half-lives, we can assign each protease’s contribution to active protein turnover. Pink, green, and purple describe the contribution of ClpP, Lon, and HslV, respectively, in stabilizing the protein. Orange marks the contribution of redundancy, defined as the additional gain in stability upon simultaneously knocking out the three proteases as compared to the additive contributions of individual KOs. Blue represents the fraction that is unexplained by the three proteases. The bar graph represents examples from each of the six categories - turnover explained predominantly by ClpP, Lon, HslV, additive contributions, redundant contributions, and actively degrading proteins in the triple KO. **E)** Bar graph for the number of substrates and the % of the proteome assigned to each of the six categories described in D. **F)** Comparison of the substrates from our categories in E with previous proteome-wide substrate-protease assignment studies, shown as overlap *p*-values. ClpP trapped substrates significantly overlap with the identified ClpP substrates (Fisher test, *p*-value = 6E-9), and previously identified substrates of HslV show a significant overlap with redundant and additive substrates (Fisher test, *p*-value = 2E-3). **G)** Comparison of the percentage of active turnover per hour across the protease KOs under N-lim. In WT cells, 4.5% of the proteome is replaced by active degradation each hour. The proteome of Δ*hslV* cells turns over at the same rate as WT cells. Both Δ*clpP* and Δ*lon* cells had 30% less protein turnover than WT cells. Even after knocking out *hslV, lon*, and *clpP* simultaneously, 40% of the WT proteome-turnover remains, suggesting that a major pathway of protein degradation in *E. coli* remains to be discovered.

Deleting *ftsH* is more complicated than the other proteases. One of its substrates, LpxC, catalyzes the committed step in the lipid A synthesis pathway. Lipid A is the hydrophobic anchor of lipopolysaccharides (LPS), a critical outer membrane component. Deletion of *ftsH* leads to increased levels of LpxC, causing an accumulation of LPS that makes the cells nonviable ^54^. *ftsH* null cells can be rescued with a mutation of FabZ (L85P), which slows LPS synthesis and compensates for the increased LpxC levels.^54^ Interestingly, we were only able to generate the ΔftsH fabZ (L85P) strain in DY378 background.^55^ Our attempts to knock out ftsH in the NCM3722 background used for the remainder of this paper failed. We are currently investigating which other modifications in DY378 might make ΔftsH fabZ (L85P) viable. These Δ*ftsH fabZ (L85P)* cells are viable, though unfortunately, they grow too slowly on minimal media and are washed out of the chemostat. Therefore, we could not measure protein turnover in a *ftsH* mutant in a similar manner to the other proteases. Instead, we repeated the batch nitrogen starvation experiments (Fig. 2E). Similar to the WT cells, cells lacking *ftsH* degraded cytoplasmic proteins. In contrast, membrane proteins are mostly stable (Fig. 3B). This indicates that none of the four known ATP-dependent proteases in *E. coli* are individually responsible for the large-scale cytoplasmic recycling that occurs under nitrogen limitation.

We then asked if proteases might act redundantly, i.e., multiple proteases share a substrate, which could mask the effects of deleting individual proteases. To this end, we measured protein turnover in a triple KO line (Δ*hslV* Δ*lon* Δ*clpP*) in nitrogen limitation. A quantitative comparison of protein degradation rates between the triple KO and the individual KOs allows us to assign the contribution of each protease in turning over a substrate (Fig. 3D). We can classify the substrates into six groups: those being degraded predominantly by a single protease, those where the effects of the individual proteases are additive, those that are stabilized more in the triple KO than the combined effect of individual KOs (redundantly degraded), and those that are still actively degraded in the triple KO (Table S2).

We classified 64 and 14 substrates to be predominantly degraded by ClpP and Lon, respectively. We only assigned one substrate uniquely to HslV: UhpA, a transcriptional regulator that activates the transcription of genes involved in transporting phosphorylated sugars.^56^ 81 proteins are degraded additively, a notable example of which is IbpA, a small chaperone. Previous studies have proposed that Lon degrades free IbpA/lbpB and bound client proteins.^57^ We found that ClpP and Lon contribute approximately equally to the degradation of IbpA, and their contribution is additive.

We classify 37 proteins as being redundantly degraded by two or more proteases. For example, YbhA is rapidly degraded in all single KO strains but stabilized in the triple KO, indicating that at least two of these proteases act redundantly. Interestingly, the majority of cytoplasmic proteins are still slowly degraded in the triple KO under nitrogen limitation, and we classified 94 proteins as still being actively degraded (Fig. 3E). LexA, an SOS repressor, auto-degrades itself under stress and unperturbed growth.^58,59^ Consistent with this, LexA still undergoes degradation in the triple KO. It will be interesting to investigate if other proteins with short half-lives in the triple KO are auto-degrading, degraded by FtsH, or if other mechanisms are at play.

To validate our classifications, we compared our protease-substrate relationships with previous proteome-wide measurements. We see a significant overlap (*p*-value = 6E-9) of our identified ClpP substrates with substrates identified via a trap mutant (Fig. 3F).^20^ However, we do not observe an overlap of our putative Lon-substrates with a previous Lon-trap experiment (*p*-value = 0.27).^21^ This lack of overlap is most likely caused by our separating the Lon trap substrates into the different classifications, indicated by a more significant overlap with the substrates that were stabilized in any of our KO strains (*p*-value = 0.05). This is consistent with previous observations that Lon substrates are often shared with other proteases.^60^ Interestingly, the putative substrates of HslV identified through a microarray study show a strong overlap with the proteins we classify as additive or redundant (*p*-value = 0.002).^23^ This is consistent with previous reports that HslV substrates are shared with other proteases.^26,61^ We also found mild enrichment (*p*-value = 0.08) between substrates identified in a previous FtsH trap^62^ study and additive or redundant substrates, consistent with findings that FtsH often degrades proteins that are also substrates for other proteases.^27^ The lack of overlap between the proteins still degrading in the triple KO and FtsH-trap substrates implies that FtsH is likely not involved in the degradation of these substrates.

Surprisingly, 40% of active protein degradation in nitrogen limitation in wild-type cells persists upon knocking out the three canonical ATP-dependent cytoplasmic proteases (Fig. 3G). We could not generate a viable quadruple KO with *ftsH* deletion, so we cannot rule out the possibility that all four proteases act redundantly as an explanation of the remaining protein degradation. However, the results from the individual *ftsH* knockout (Fig. 3B) and the lack of overlap between degrading proteins and the FtsH-trap experiment (Fig. 3F) are evidence against FtsH being responsible for the remaining degradation. Regardless, a major pathway for degrading proteins in *E. coli* remains to be discovered: either FtsH plays a much bigger role than is currently believed, or a completely new mechanism degrades cytoplasmic proteins under nitrogen starvation.

### Analyzing features of rapidly turning over proteins

We found that most short-lived proteins have similar half-lives regardless of nutrient limitation (Fig. 4A, Table S3). With gene-set enrichment,^63^ we found that rapidly degraded proteins were enriched in transcriptional regulators (Benjamini-Hochberg adjusted *p*-value 4E-4). A protein’s response time depends on its replacement rate.^5^ Proteins involved in transcriptional regulation might need to rapidly adjust their levels to changing growth conditions.

**Figure 4:**
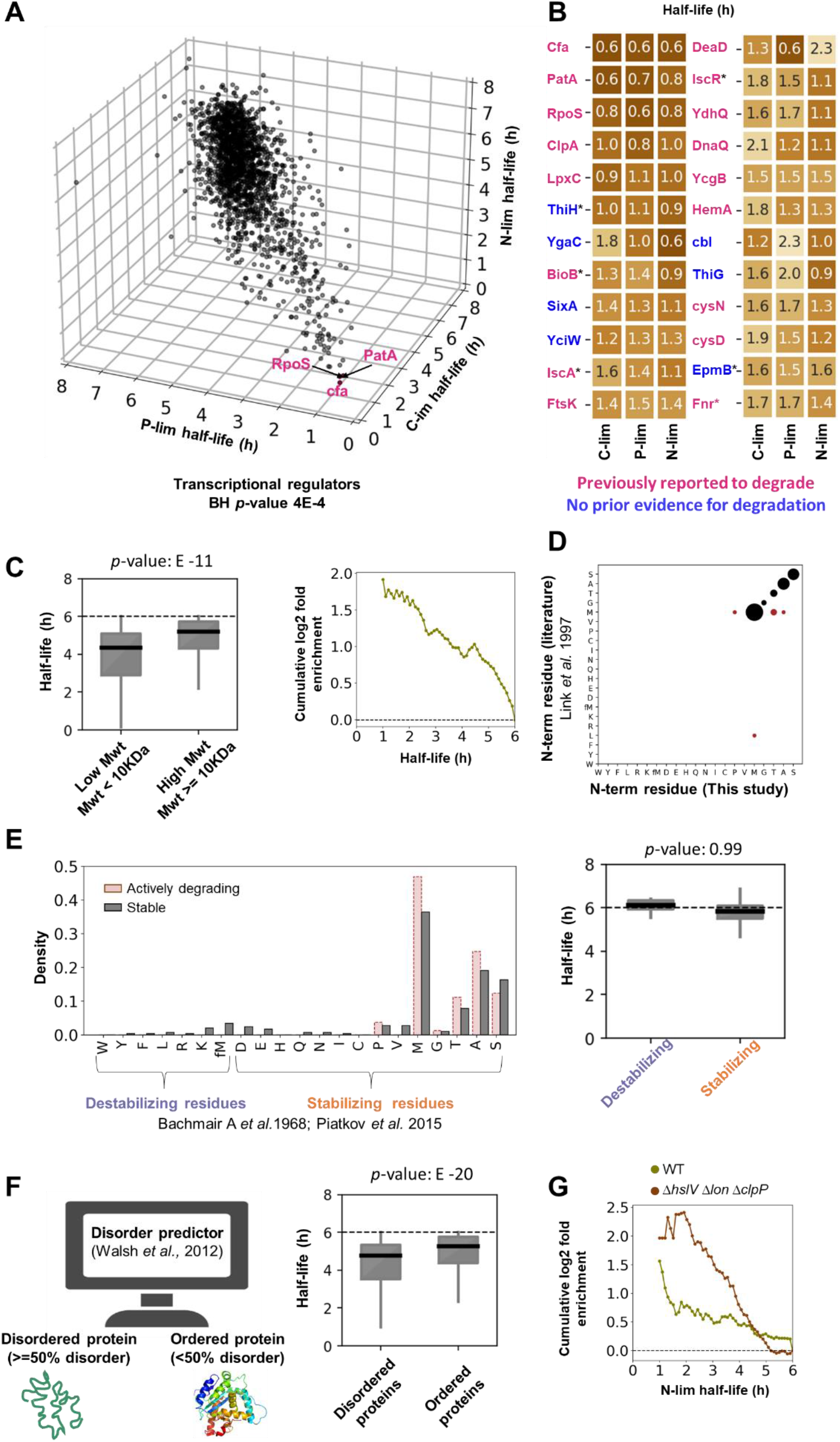
Features of proteins with short half-lives. **A)** Scatter plot of protein half-lives for C-lim, P-lim, and N-lim conditions. The half-lives of rapidly degrading proteins are typically similar under different nutrient limitations. Labeled are the three proteins with the shortest average half-lives. Short-lived proteins are enriched for transcriptional regulators (BH *p*-value 4E-4). **B)** The 24 proteins with the shortest mean half-lives in the three nutrient limitations. For eight of these proteins (in blue), we could not find any prior literature evidence for degradation, and six (marked with *) contain Fe-S clusters (BH *p*-value = 0.048). **C)** Smaller proteins tend to have shorter half-lives. Left: Box plot of half-lives averaged over all the nutrient limitations for low and high molecular weight proteins. Smaller proteins have significantly lower half-lives than larger proteins (*p*-value: E-11). Right: Fold enrichment of low versus high Mwt proteins as a function of half-life. Rapidly turning over proteins are enriched for the smaller proteins. **D)** Comparison of the proteome wide N-terminus amino acid residues obtained from this study and prior literature. The size of the marker indicates the number of proteins with a particular residue. **E)** We see no correlation between the N-terminal protein residue and protein half-lives (N-end rule). Left: Distribution of the N-terminus residues for actively degrading and stable proteins. The canonical destabilizing N-terminus amino acids show no enrichment for actively degrading proteins. Right: Box plots of half-lives for the canonical destabilizing and stabilizing residues indicate no detectable separation in stability based on their N-terminus (*p*-value: 0.99). **F)** Disordered proteins tend to have shorter half-lives. Left: The Espritz algorithm classified proteins as ordered or disordered. Right: Box plot of half-lives for disordered and ordered proteins. Disordered proteins have significantly lower half-lives than ordered proteins (*p*-value: E-18). **G**) Fold enrichment of disordered versus ordered proteins as a function of half-life for the WT (olive) and triple protease KO (brown) cells. Rapidly turning over proteins are enriched for the disordered category. This enrichment becomes more pronounced when three proteases were knocked out, suggesting that many of the remaining short-lived proteins could be degraded in an ATP-independent manner.

Using our data set, we validated examples of degradational regulation that had previously been reported and also uncovered novel targets. Of the 24 proteins with the fastest average degradation rates, 17 were previously reported to be degraded (Fig. 4B). Seven proteins—ThiH, YgaC, SixA, YciW, CbI, ThiG, EpmB—had no prior evidence in the literature for degradation. Interestingly, six rapidly degrading proteins—ThiH, BioB, IscA, IscR, EpmB, Fnr—contain Fe-S clusters, which is significantly higher than expected by random chance (BH *p*-value = 0.048). Flynn et al. previously proposed that Fe-S binding proteins are degraded under aerobic conditions, likely because the Fe-S clusters are oxidized, destabilizing the protein.^20^.

Multiple metabolic enzymes such as PatA, LpxC, and HemA are also rapidly degraded. Rapid degradation allows for immediate and direct control over intracellular protein levels based on cellular demand. PatA (Putrescine-Aminotransferase) is involved in putrescine (polyamine) degradation (*K*_M_ = 9 mM) and is unstable under standard growth conditions with high putrescine levels, in which another enzyme (PuuA - glutamate-putrescine ligase) dominates usage of putrescine. PatA is expected to stabilize in specific growth conditions with low putrescine concentrations.^64^ LpxC, a protein required for lipid A synthesis, is rapidly degraded under slower growth to balance LPS production with cellular demand.^54^ HemA, involved in porphyrin biosynthesis, is degraded when the media lacks heme as an iron source.^65^ DnaQ, the proofreading exonuclease of the stable DNA polymerase III core enzyme [DnaE][DnaQ][HolE], is rapidly degraded (*t*_1/2_= 1.2 h). DnaE is more stable with a half-life of 5 h, whereas HolE is undetected in our data set, likely due to its short length. Free DnaQ is unstable but stabilized on complexation with HolE.^66^

We next tested whether these rapidly degrading proteins share attributes such as their physiochemical properties, sequence features, or structural characteristics. We found that smaller proteins (MW < 10 kDa) have significantly shorter half-lives regardless of the nutrient limitation (Fig. 4C, Fig. S2A). This enrichment was more pronounced at lower half-life cutoffs (Fig. 4C). On the other hand, charge and isoelectric point were not significantly correlated with half-lives under P-lim and C-lim (Fig. S2B, C). However, both the charge and the isoelectric point of a protein were correlated with half-lives under N-lim (Fig. S2B, C *p*-value= E-20). This is most likely because cytoplasmic proteins are short-lived under N-lim while membrane proteins are typically stable. Membrane proteins tend to have higher isoelectric points and more positive charge due to their interaction with negatively charged phospholipids.^67,68^

One obvious sequence feature to investigate is the N-end rule, which relates a protein’s stability to its amino-terminal residue.^69^ Amino-terminal arginine, lysine, leucine, phenylalanine, tyrosine, tryptophan, and formylated N-terminal methionine (fMet) are believed to be destabilizing residues, whereas the other residues are believed to be stabilizing.^69,70^ First, we determined the *in vivo* N-terminal residue of ∼600 proteins using a label-free proteomics data set (Table S4).

MS2 spectra were searched with the Sequest algorithm^71^ considering all possible N-terminal tryptic subfragments for a protein. Encouragingly, when we compared a small subset of the identified N-termini with previous data in literature obtained using Edman Degradation,^72^ we found nearly perfect agreement (55/61 proteins)(Fig. 4D). Surprisingly, however, our rapidly degrading proteins showed no enrichment for the previously reported destabilizing N-terminal amino residues (Fig. 4E). This suggests that the N-terminus of *E. coli* proteins is not the primary determinant of proteins’ *in vivo* stability.

Another sequence feature previously shown to affect protein stability in bacterial and eukaryotic cells is intrinsically disordered protein segments.^73,74^ To this end, we determined the percentage disorder for all the proteins using the Espritz algorithm.^75^ Disordered proteins had significantly shorter half-lives than ordered proteins (Fig. 4F, *p*-value: E-208). Interestingly, this enrichment further increases when we use protein half-lives measured in the triple protease knockout cells (Δ*hslV* Δ*lon* Δ*clpP*) (Fig. 4G). This is consistent with ATP-dependent proteases being able to unfold and digest structured proteins. Once these proteases are removed, the remaining proteins with short half-lives should be enriched for those that are unstructured and therefore prone to degradation by energy-independent proteases.

### Analysis of turnover for functionally related proteins

Next, we investigated protein turnover for functionally related protein modules, such as multiprotein complexes, operons, and metabolic pathways. We calculated each module’s coefficient of variation (CV) and compared this to the CV distribution when proteins were randomly assigned to sets. We observe that the functionally associated modules exhibit significantly lower variance than if the proteins were randomly assigned to each module (Fig. 5A), suggesting that functionally associated proteins tend to exhibit similar half-lives.

**Figure 5:**
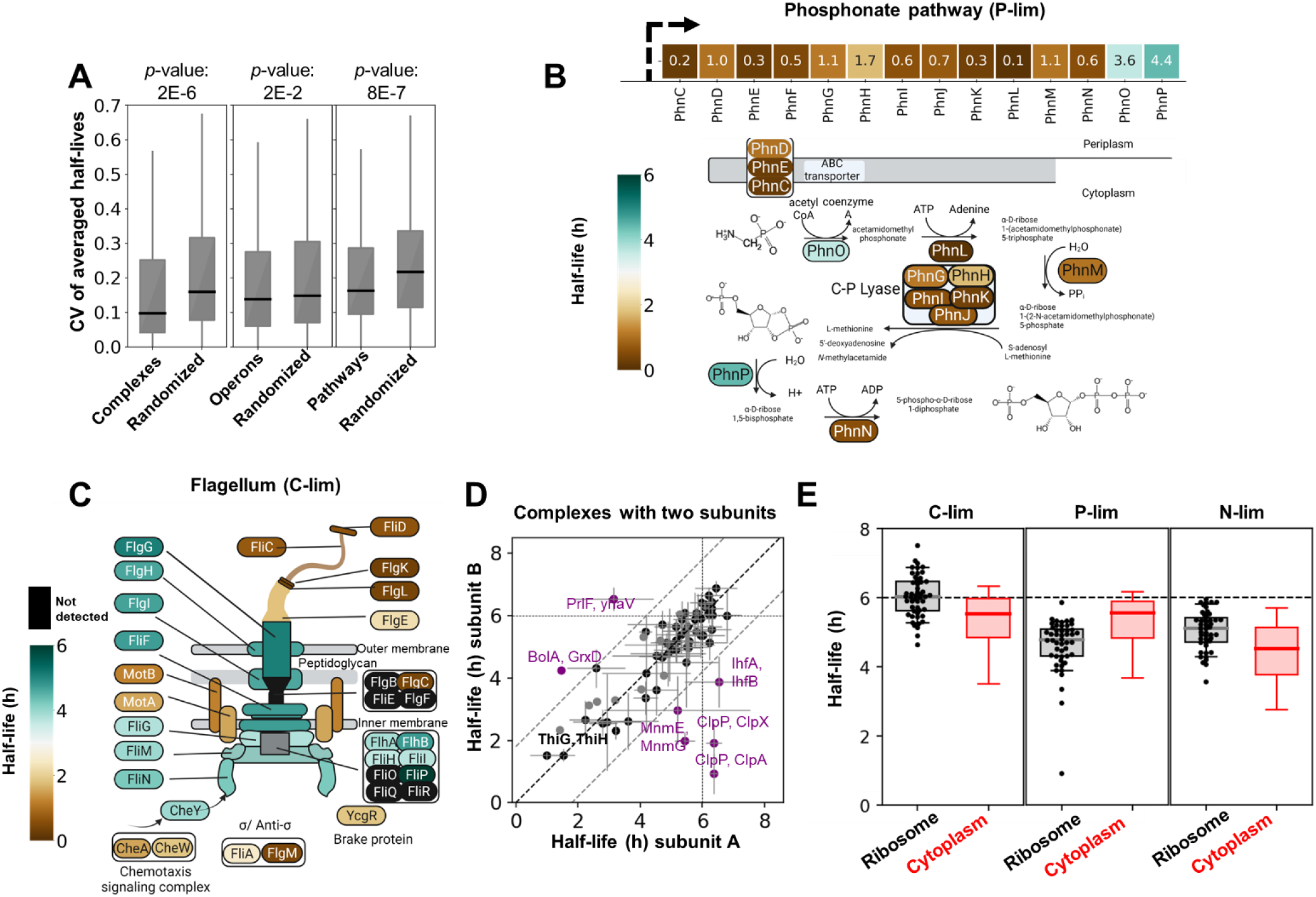
Protein half-lives of functionally related proteins. **A)** Functionally related proteins tend to have similar half-lives. Boxplots of CVs of half-lives for functionally related proteins compared with a randomized set of proteins (shown *p*-values are Wilcoxon rank sum test). **B)** Map of the half-lives for proteins involved in the phosphonate pathway under P-lim. The phosphonate pathway is encoded in a single operon and is involved in the transport and metabolism of organophosphonates (C-P bond). Most of these proteins degrade rapidly under P-lim. **C)** Map of the half-lives of flagellar proteins under C-lim. The basal body elements of the flagellum are largely stable, while filament and motor structure components are rapidly degrading. **D)** Scatter plot of protein half-lives for protein complexes containing two distinct subunits. Error bars represent standard deviation. Highlighted in purple are complexes for which the half-lives of subunits disagree, potentially because their interaction is transient, e.g., ClpA/P or BolA, GrxD. **E)** Box plots of half-lives for ribosomal proteins compared to all cytoplasmic proteins. Ribosomal proteins are more stable than cytoplasmic proteins in C-lim and N-lim but degrade faster than the median of cytoplasmic proteins in P-lim.

For example, 12 of the 14 proteins involved in phosphonate metabolism and transport are rapidly degraded (average half-life = 0.7 h) under P-lim (Fig. 5B). These proteins were 16-fold more abundant in P-lim compared to N-lim and C-lim (Table S5). Therefore, we were unable to measure their half-lives in C-lim or N-lim, so it’s unclear if they turn over in these limitations as well. Additionally, proteins associated with flagella show correlated expression levels and half-lives. Surprisingly, most of the proteins forming the basal flagellar body are stable, but the filament (FliC, FliD), motor (MotA, MotB), and sensory proteins (CheA, CheW) are rapidly degraded (Fig. 5C). Future work will be required to decipher the underlying mechanisms and functional relevance.

In general, proteins that form a complex tend to exhibit similar half-lives (Fig. 5D). Several complexes whose subunits are degraded at different rates are known to interact weakly or transiently or have subunits which are expressed non-stoichiometrically, suggesting that at least some of these discrepancies might be due to annotation details. For example, ClpA and ClpX are the unfoldases in complex with ClpP. Autodegradation of ClpA is used to regulate the number of ClpAP complexes in the cell and the flow of substrates to ClpAP.^52^ Finally, antitoxins like PrlF are subject to regulated degradation while their toxin counterparts are stable.^76,77^

The ribosome is one of the heterocomplexes which exhibits unanticipated patterns under different nutrient limitations (Fig. 5E). Under both C-lim and N-lim, ribosomal proteins are slightly more stable than the median cytoplasmic protein. However, under P-lim, ribosomal proteins are less stable than typical cytoplasmic proteins. rRNA contains about 50% of the cellular phosphorus.^78^ Therefore, cells likely recycle the phosphorus stored in rRNA when phosphorus is scarce, and associated ribosomal proteins might become unstable once their binding partners are lost.

### Active protein degradation rates typically do not scale with division rates

So far, we have compared degradation rates under various nutrient limitations but with the same cell division time. We wanted to determine how protein degradation scales with cell cycle time. The total turnover rate (*k*_total_) of a protein is a combination of active degradation (*k*_active_) and dilution (*k*_dilution_) due to cell division (Fig. 6A). We consider two simple and reasonable models of the relationship between these two parameters. In the first model, *k*_active_ scales with *k*_dilution_, i.e., the protein half-life remains a constant fraction of the cell cycle time. In the second model, active degradation rates are independent of growth rate, i.e., the active degradation rate of each protein remains constant regardless of cell doubling time.

**Figure 6:**
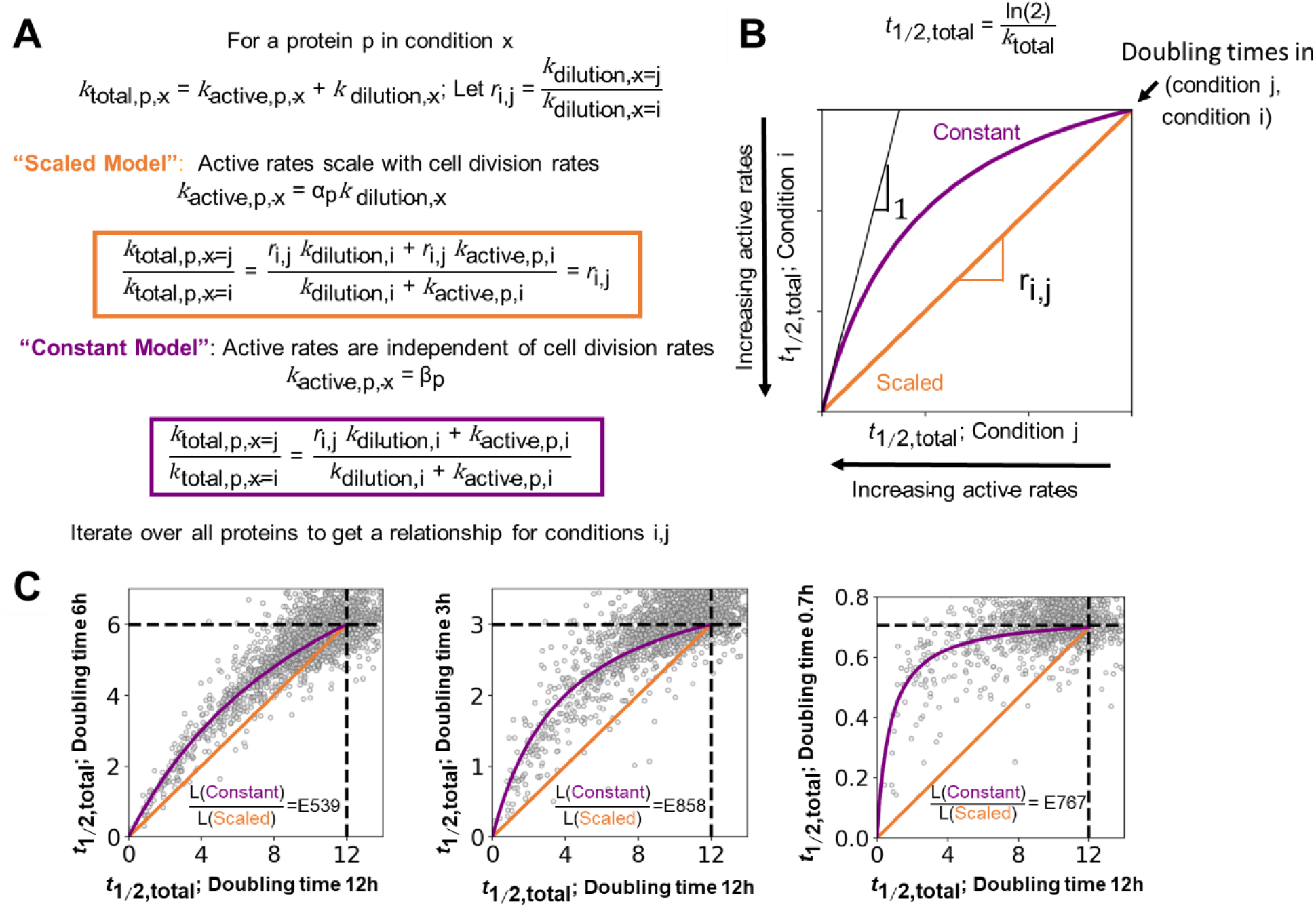
Active degradation rates are generally uncoupled from cell division rates. **A)** Two simple models describing the relationship between cell division and protein-specific turnover rates. The total protein turnover rate (ktotal) is the sum of the active degradation rate (k_active_) and the dilution rate due to cell division (k_dilution_)In the “scaled model,” active degradation rates increase proportionally to division rates with a protein-specific constant (α_p_), i.e., active degradation remains a constant fraction of the total protein turnover rate. In the “constant model,” protein-specific active degradation rates are constant (β_p_), regardless of changing division rates. In this case, for slower-dividing cells, the contribution of active degradation increases relative to dilution. **B)** *t*_1/2, total_ is the time taken to replace half the protein. A theoretical plot of *t*_1/2, total_ from two conditions (i, j) where cell division rates change by a factor of r < 1. In the scaled model, *t*_1/2, total_ values for all the proteins lie on a straight line with slope r (orange). In the constant model, the *t*_1/2, total_ values follow a nonlinear relationship between the two doubling times (purple). For proteins with very high active degradation rates, the constant model predicts that *t*_1/2, total_ will approach the same value for both doubling times, indicated by the slope 1 line (black). For diluting proteins with no active degradation, both models converge to the doubling times of conditions i and j. **C)** Scatter plots of protein *t*_1/2, total_ for *E. coli* grown at doubling times of 6 h (C-lim), 3 h (C-lim), and 0.7 h (defined minimal media batch) compared to 12 h (C-lim). The dotted lines represent the dilution limit. We observe a strong statistical preference for the “constant model,” in which active degradation rates are uncoupled from cell cycle duration. Shown are the likelihood ratios (L) of the constant models compared to the scaled models assuming normally distributed errors.

The two models have distinct predictions on how the total protein replacement half-time (*t*_1/2, total_) should scale with changing cell cycle times. In the scaled model, *t*_1/2, total_ for each protein linearly increases with cell cycle time (Fig. 6B). In contrast, in the constant model, the dilution rate dominates for rapidly dividing cells while the contribution from active degradation becomes more relevant for slower dividing cells.

To test the models’ predictions, we grew *E. coli* cells with a range of doubling times, including rapidly doubling cells with unlimited growth in minimal medium (0.7h) and slower doubling cells in carbon-limited chemostats (3h, 6h, and 12h). We found that the data favored the constant model regardless of which cell-cycle times we compared (Fig. 6C). This indicates that active protein degradation rates typically remain constant regardless of cell division rates.

We noted interesting exceptions to the model, however, particularly when comparing slower-growing cells in the chemostat to cells growing without nutrient limitation. For example, RpoS degrades faster in unlimited growth conditions than the non-scaled model would predict based on chemostat measurements. This is consistent with the previous finding that RpoS is rapidly degraded in exponentially growing cells but becomes stabilized when nutrient-limited.^79^

Because *k*_active_ rates are generally constant across cell division rates, we can more accurately measure *k*_active_ when *k*_dilution_ is small. Importantly, the observed constancy of degradation rates, regardless of cell cycle times, allows us to extrapolate active degradation rates from conditions with long cell division times (e.g., chemostats with 6-hour doubling times) to conditions with more rapid cell division times, in which separating between active degradation rates and dilution rates is experimentally difficult. Thus, the protein half-lives in this manuscript, primarily obtained in the chemostat, are a valuable resource that can be extrapolated to arbitrary cell division rates.

## Discussion

This paper introduces a technique for the global measurement of protein turnover on a gene-by-gene basis by combining complement reporter ion quantification with heavy isotope labeling of nutrients (Fig. 1). Applying our method to measure protein turnover across multiple nutrient limitations, we found that most cytoplasmic proteins slowly degrade in nitrogen-limited conditions (Fig. 2). By contrast, in phosphorus-limited and carbon-limited conditions, proteins are mostly stable. We observe this phenomenon in a nitrogen-limited chemostat and in a nitrogen-starved batch culture. The slow degradation of cytoplasmic proteins is likely a strategy

*E. coli* has developed to keep scarce amino acids available, which could be critical to various metabolic processes, including the ability to synthesize new proteins and adapt the proteome to changing environments. Bulk protein turnover measurements in the 1950s showed that *Saccharomyces cerevisiae* also increases overall protein turnover when starved of nitrogen,^15^ suggesting that a similar strategy might apply to eukaryotes.

We assigned protein substrates to proteases by measuring the change in protein turnover rates in protease knockout strains (Fig. 3). We were surprised by how little protein degradation changed in the knockout strains, particularly when deleting the canonical proteases ClpP and Lon. We showed that in these knockout strains, only a few proteins have a slower degradation rate, and the observed degradation of cytoplasmic proteins continues. Even when we knocked out *clpP, lon*, and *hslV* simultaneously, 40% of total protein turnover remains, including the cytoplasmic recycling and the degradation of many short-lived proteins. However, we observe remarkable additive and redundant effects when comparing protein turnover rates in the individual knockouts with the triple knockout. This suggests that many proteins are substrates for more than one protease.

We could not extend these approaches to identify substrates for FtsH, as its deletion is lethal due to the accumulation of LPS. However, when combined with a *fabZ* mutation,^54^ we could show that the degradation in nitrogen-starved batch culture continues when *ftsH* is deleted. The lack of significant overlap between proteins that are still degraded in the Δ*clpP*Δ*lon*Δ*hslV* strain and proteins pulled down from an FtsH-trap (Arends et al. 2016) also suggests that FtsH is likely not responsible for the remaining degradation. So far, we have not been able to generate viable quadruple knockout cells for all four known ATP-dependent proteases in *E. coli*. We, therefore, cannot completely rule out that FtsH is responsible for the remaining cytoplasmic degradation when the other three proteases are deleted. Regardless, a major protein degradation pathway in *E. coli* still needs to be discovered: either FtsH plays a much more significant role than generally anticipated, or there is an entirely different pathway outside the four known ATP-dependent proteases. While protein degradation itself is energetically favorable, unfolding a protein requires energy. *E. coli* encodes many non-ATP-dependent proteases,^80^ but the rapid turnover of proteins in the triple knockout line and the fact that most cytoplasmic proteins are structured suggest that some ATP-dependent unfoldase is involved. Perhaps, an adapter like ClpX or a chaperone unfolds proteins and allows those substrates to be degraded by one of the proteases believed to be energy-independent.^80^

We found that many proteins have short half-lives regardless of nutrient limitations (Fig. 4). Among those, we see an overrepresentation of transcriptional regulators. Rapid turnover might enable a quick response to changing growth conditions to rapidly adjust transcription rates to the new environment. Surprisingly, we found no correlation between a protein’s half-life and its N-terminal residue suggesting that the N-end rule is a poor predictor for protein stability *in vivo*. In contrast, disordered proteins are drastically enriched among proteins with rapid turnover (Fig. 4F). This might suggest that many short-lived proteins could be degraded in an energy-independent way. This is further supported by our finding that the enrichment of disordered proteins is further increased among proteins that are degraded when the three ATP-dependent proteases ClpP, Lon, and HslV are deleted. We observe highly significant correlation among protein turnover for functionally related proteins like complexes and those expressed from operons. We found striking examples of this regulation in phosphonate metabolism and flagella but can currently only speculate about the underlying regulatory mechanisms and functional importance. Further studies will be required to follow up on these intriguing observations.

When we compared protein turnover across *E. coli* growth rates, we found that rates of active protein degradation remain constant (Fig. 6). Based on this finding, the relative contribution of active degradation compared to dilution due to cell division must change as a function of growth rate. Therefore, relative protein levels of actively degrading proteins must change with differing cell growth rates, or cells must compensate by adjusting transcription and/or translation rates. Protein expression regulation combines gene-specific effects with such global parameter changes.^16^ Our insight will help to improve genome-wide protein abundance regulation models and could help to better engineer gene expression circuits with desired properties.

The discoveries in this manuscript have been enabled by introducing a new method to measure protein turnover. We chose to use heavy ammonia to label newly synthesized proteins to avoid the use of mutants and to boost the signal for short labeling times. The resulting MS1 and MS2 spectra are extremely complex, making standard quantification approaches challenging.^42^ We have overcome these challenges by taking advantage of the outstanding ability of the complement reporter ion strategy (TMTproC) to distinguish signals from the chemical background.^46^ The introduced methods are applicable widely beyond *E. coli*. Our ability to use comparatively cheap heavy isotope labels opens up the possibility of performing similar studies on larger animals, e.g., after D_2_O intake, which would be cost-prohibitive with heavy amino acid labeling.^42,81^ Unlike many other cutting-edge multiplexed proteomics approaches, the applied technology is compatible with comparatively simple and widely distributed instrumentation, such as quadrupole-Orbitrap instruments, as we avoid the need for an additional gas-phase isolation step. The required analysis software is available on our lab’s GitHub site (https://github.com/wuhrlab/TMTProC).

We have generated a broad resource of protein turnover rates in 12 different growth conditions, each with biological replicates. The investigated conditions include varying cell cycle times from 40 minutes to 12 hours, nitrogen-, carbon-, phosphorus-limitation, and various protease knockout strains. We expect this resource to allow researchers to complement their data sets with protein turnover information. Our finding that active degradation rates are typically constant regardless of division rates will allow researchers to extrapolate protein half-lives to arbitrary conditions. Our measurements of how protein turnover rates change in protease knockout strains will help refine protease-substrate relationships. Unlike studies relying on trap experiments or protein microarrays, we could start to deduce the redundant nature of these connections. We have shown the power of the provided resource by demonstrating cytoplasmic recycling in nitrogen limitation and by finding a scaling law for active protein degradation rates with varying cell cycle times. Thus, we advance protein turnover measurement technology, provide a resource for ∼3.2k *E. coli* protein half-lives under various conditions, and provide fundamental insight into global protein expression regulation strategies.

## Acknowledgements

We would like to thank Markus Basan for the gift of the triple-protease knockout strain. We thank Michaela Eickhoff, Yihui Shen, Josh Rabinowitz, Jonathon O’Brien, Joe Sheehan, and Irina Mikheyeva-Bridges for their advice and discussions. This work was supported by NIH grants R35GM128813 (MW), R35GM118024 (TJS), and grant T32-GM007388 (to Princeton University [EMH]), the U.S. Department of Energy, Office of Science, Office of Biological and Environmental Research under award number DE-SC0018420 and DOE grant DE-SC0018260. This work was supported in part by the National Science Foundation through the Center for the Physics of Biological Function (PHY-1734030 [NSW]). We gratefully acknowledge support by the American Heart Association predoctoral fellowship 20PRE35220061 (TN), Princeton Catalysis Initiative (MW), Eric and Wendy Schmidt Transformative Technology Fund (MW), Harold W. Dodds Fellowship (MG), NSF Graduate Research Fellowship (ERC), Princeton University’s Summer Undergraduate Research Program (EC).

## Declaration of interests

The authors declare no competing interests.

## Notes

### Competing Interest Statement

The authors have declared no competing interest.

### Summary of Updates

We have updated the manuscript and added additional figures.

