## Supplemental material for "Global Protein-Turnover Quantification in *Escherichia coli* Reveals Cytoplasmic Recycling under Nitrogen Limitation"

### Supplementary Figures

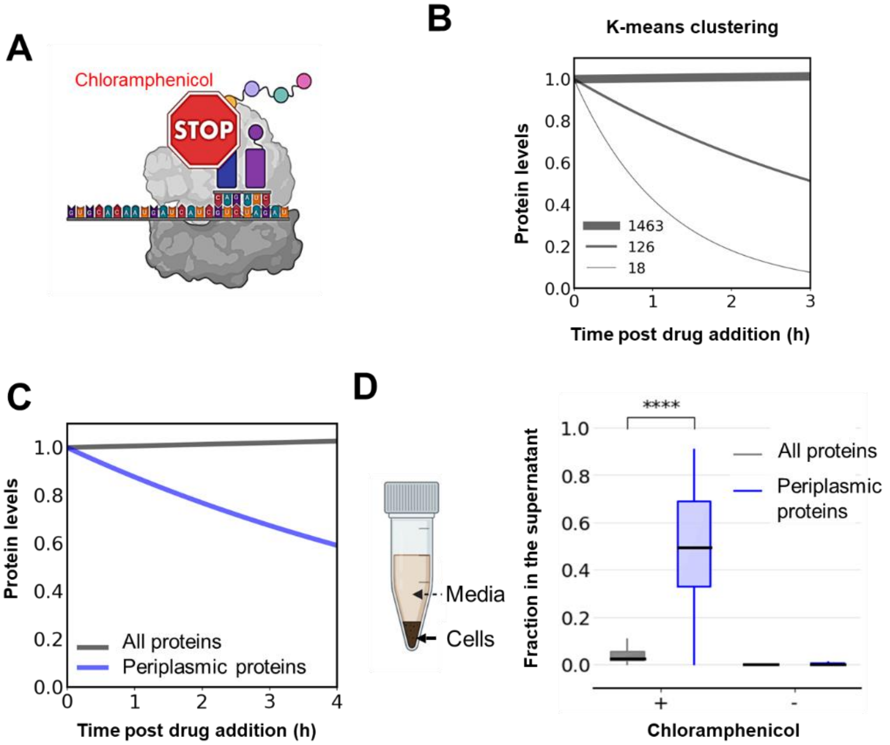


**Figure S1: Using translation inhibition to measure protein turnover might lead to unintended side effects. A)** Protein synthesis is halted by adding chloramphenicol to a culture of exponentially dividing E. coli cells. Proteomic analysis samples are collected post-drug addition, with time points fitted to an exponential decay curve using half-life as a parameter. **B-C)** The shown curves are obtained using the fitted half-lives. **B)** k-means clustered protein abundance profiles after the addition of drug. The largest cluster of proteins do not change abundance, indicating that they are stable. **C)** Median protein levels over time for all proteins and periplasmic proteins. The median periplasmic protein appears to be degrading, while most of the proteins are stable. **D)** We separated the supernatant from the cells using centrifugation and quantified relative protein levels in both samples with proteomics. We calculated the fraction in the supernatant for each protein by dividing the abundance in the supernatant by the total abundance for that protein. Our data indicate that the supernatant is enriched with periplasmic proteins following chloramphenicol addition, suggesting that chloramphenicol may prompt the release of periplasmic proteins into the supernatant.

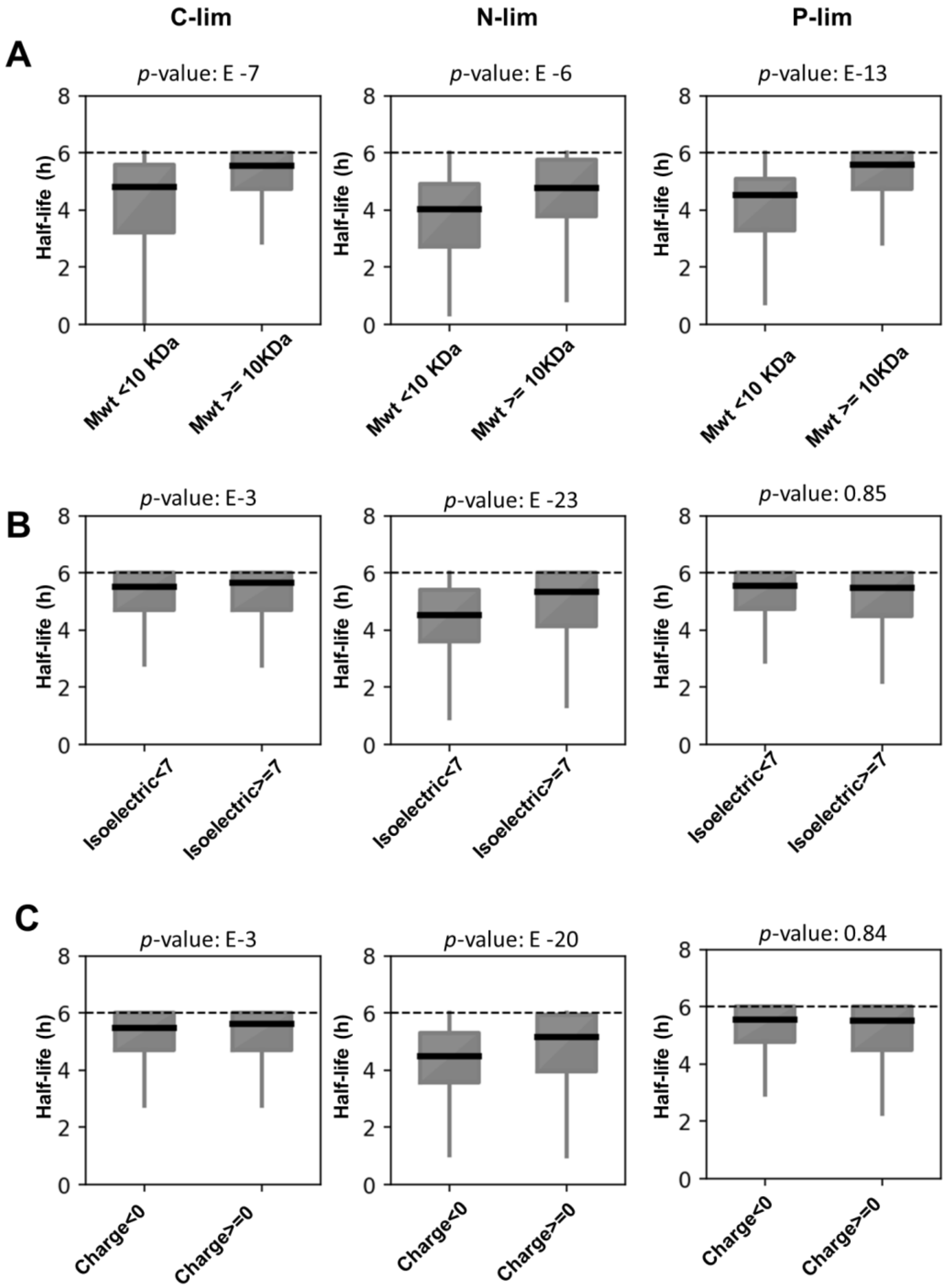


**Figure S2: Relationship between protein half-life and their physiochemical properties. A)** Box plots of protein half-lives categorized by molecular weight for C, N, and P-lim conditions. Notably, smaller proteins exhibit significantly shorter half-lives. **B)** Box plots of protein half-lives categorized by isoelectric points for C, N, and P-lim conditions. Under N-lim, shorter-lived proteins are significantly more acidic, while no significant effect is observed under C and P-lim. **C)** Box plots for half-lives of proteins separated by their charge for C, N, and P-lim. Shorter lived proteins tend to be negatively charged under N-lim, However, there is no significant effect under C and P-lim. (p-values are obtained using a one sided Mann-Whitney U rank test)

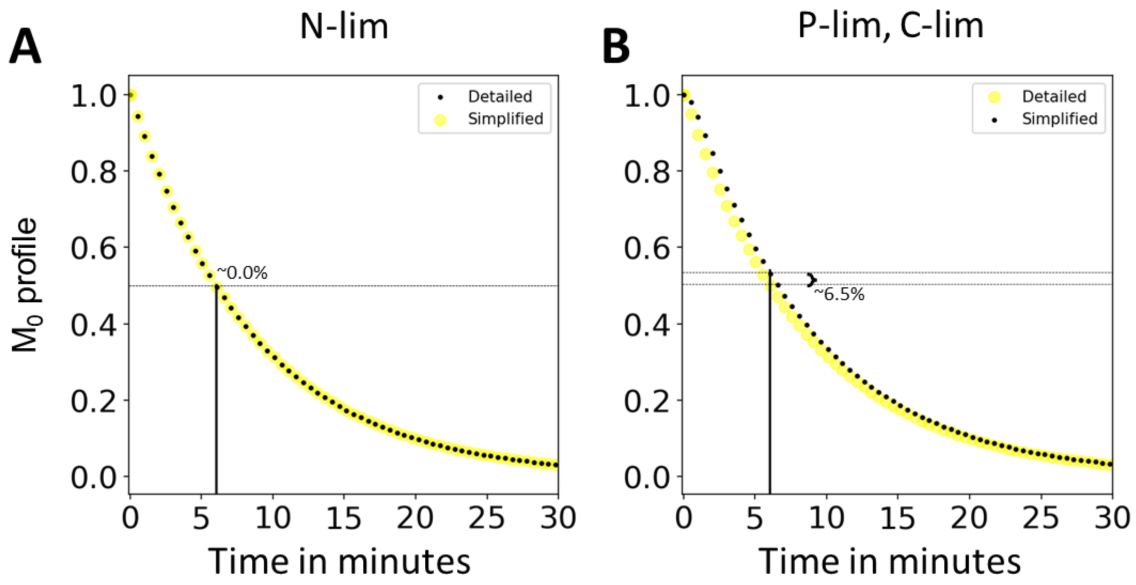


**Figure S3**: **Equivalence of the detailed and simplified model of the monoisotopic decay profile over time.** A) N-limited B). P-limited cells

### Experimental Protocols

#### Turnover measurements

*E. coli* strain NCM3722 was grown in continuous culture at 37 °C. The cells were cultured in chemostats as described previously (Citation). The chemostat (Sixfors, HT) volume was 300 mL with oxygen and pH probes to monitor the culture. pH was maintained at 7.2 ± 0.1. 40 mM MOPS media (M2120, Teknova) was used with glucose (0.4% w/v, Sigma G8270), ammonia (9.5 mM $NH_4Cl$, Sigma A9434) and phosphate (1.32 mM $K_2HPO_4$, Sigma P3786) added separately. For C- and N-limiting media, glucose and ammonia concentrations were reduced by 5-fold (0.08% and 1.9 mM, respectively). The P-limiting medium contains 0.132 mM $K_2HPO_4$. Cells were grown up to steady state in a media containing light ammonia for ~ 8-10 generations. This is equivalent to running the 6 h doubling time chemostat for ~ 2.5 days before switching the feed to a media containing heavy ammonia (CIL, NLM-107-10-10). Based on the $OD_{600}$, ~4-6 mL of effluent is collected after the switch for proteomics analysis. The $OD_{600}$ for the chemostats are as follows: P-lim OD ~0.71, N-lim OD ~0.51, C-lim OD ~0.68. The effluent is immediately frozen in liquid nitrogen.

For the minimal media no limitation batch cultures, NCM3722 cells were grown at 37 °C in 40 mM MOPS media with glucose (0.4%), ammonia (9.5 mM) and phosphate (1.32 mM) added separately. The cells are grown overnight in the medium containing light nitrogen and are diluted 500x in the fresh media containing light ammonia. The cells are grown up to an OD ~0.4 before diluting by 10x into medium containing heavy nitrogen. We used $^{15}$N-ammonia rather than labeled amino acids because of the higher signal we obtain with a low labeling fraction. For example, when 10% of the nutrient pool is labeled: If using heavy arginine, 90% of newly synthesized tryptic peptides ending in arginine are unlabeled. In contrast, when using heavy ammonia, a peptide having 15 nitrogens, only $(90\%)^{15}$=21% will be unlabeled. Furthermore, we would have to work with auxotroph mutants when using amino acid labeling. Subsequently, samples of ~200 μg of protein per time point were collected. To maintain the cells in steady state, they are repeatedly diluted into fresh media after it reaches the OD~ 0.4.

The time points for proteomics collection depend on the doubling time of the bacteria. The following table details the time point used for the sample collection for all doubling times analyzed in this study:

| **Doubling time** | **Reactor type** | **Time points for sample collection (mins)** |
|---|---|---|
| 42 mins | Batch | 0, 5, 12, 20, 30, 45, 60,175 |
| 3 h | Chemostat | 0, 45, 105, 186, 270, 366, 510, 729 |
| 6 h | Chemostat | 0, 107, 229, 376, 548, 735, 1024, 1612 |
| 12 h | Chemostat | 0, 173, 302, 444, 649, 873, 1230, 2166 |

#### Chloramphenicol translation inhibition assay

*E. coli* strain NCM3722 was grown in batch cultures at 37 °C. 40 mM MOPS media (M2120, Teknova) was used with glucose (0.4%, Sigma G8270), L- arginine (2.37 mM, Sigma A5006) and phosphate (1.32 mM $K_2HPO_4$, Sigma P3786) added separately. The doubling time was 2 hours. As the cells reached exponential growth, OD~0.3, translation inhibition drugs were added to arrest the protein synthesis. Chloramphenicol was added at 200 mg/mL. *E. coli* cells (OD: 0.3 4 mL culture gave ~ 200 μg of protein) were harvested by centrifuging at 5000 g for ~ 2 mins. 10 samples were collected post the addition of drugs at [0,10,20,40,60,80,100,120,180, 240] minutes for the proteomic analysis.

#### Batch starvation assay

An overnight culture of *E. coli* grown in Luria-Bertani (LB) broth was diluted 1:100 in minimal media (40 mM MOPS media (M2120, Teknova), 0.4% glucose (Sigma G8270), 9.5 mM ammonium chloride (Sigma A9434), and 1.32 mM potassium phosphate dibasic (Sigma P3786)). Cultures were shaken at 30 °C until an OD of ~0.5 was reached. The culture was collected and centrifuged at 5 krcf for 10 minutes. The supernatant was discarded, and cells were washed three times with 40 mM MOPS, centrifuging between each wash. Cells were then re-suspended in identical minimal media but lacking either glucose or ammonium chloride. The culture was then shaken at 30 °C and samples were taken at the following times (in minutes): 0, 151, 226, 407, 408, 944. The samples were prepared for proteomic analysis using the protocol described in Sample Preparation.

#### Strain construction

The *ΔclpP*, *Δlon*, *ΔhslV* single mutants were generated by P1 transduction from the Keio collection[1] into *E. coli* strain NCM3722. The *ΔclpPΔlonΔhslV* triple knockout was provided by the Basan lab.[2]

The *ΔftsH* strain was generated as follow. First, *fabZ*$_{L85P}$ was moved into the λ Red strain DY378 with *cadR*-IG-*yaeH*::*cat* by linkage transduction.[3] Transductants were selected for on LB supplemented with 20 μg mL$^{-1}$ chloramphenicol and screened for the *fabZ*$_{L85P}$ mutation by DNA sequencing (Genewiz, South Plainfield, NJ). The *ftsH* coding sequence in the *fabZ*$_{L85P}$ mutant was deleted and replaced with the kanamycin resistance cassette and flanking flippase recognition target sites from pKD4 using l Red-mediated recombination.[4]

*Strains and plasmids*

| **Name** | **Description** | **Source or reference** |
|---|---|---|
| *Strains* | | |
| DY378 | W3110 λ*c*I*857* Δ*(cro-bioA)* | [4] |
| RLG986 | DY378 *fabZ*$_{L85P}$ *cdaR*-IG-*yaeH*::*cat*; Cam$^R$ | This study and [5,6] |
| RLG1017 | DY378 *fabZ*$_{L85P}$ *cdaR*-IG-*yaeH*::*cat* Δ*ftsH*::*kan*; Cam$^R$ Kan$^R$ | This study and [5,6] |
| *Plasmids* | | |
| pKD4 | Template plasmid containing a FRT-flanked kanamycin resistance cassette; Amp$^R$ Kan$^R$ | [7] |

*Primers*

| **Name** | **Sequence (5' – 3')** |
|---|---|
| ftsH.Fwd | GCGCTAGAAATGTGTCGTGA |
| ftsH.Rev | GGATGGCTAAGGTCCAGTGA |
| KOftsHpKD4.Fwd | CGCTGTTTTTAACACAGTTGTAATAAGAGGTTAATCCCTTGAGTGAC atgTGTGTAGGCTGGAGCTGCTTC |
| KOftsHpKD4.Rev | GTACAAATACAGTCATCTGATGCGGGAACttaCTTGTCGCCTAACTGC TCATGGGAATTAGCCATGGTCC |
| fabZ.Fwd | CTGAACCAGGCGTCTATTCC |
| fabZ.Rev | GACATGGGGTCCAACGATAC |

#### Sample preparation and data analysis for quantitative proteomics

Samples were mostly prepared as previously described.[8] Each sample containing ~200 μg of total protein was lyophilized to remove the water and then resuspended in 200 μL of lysis buffer containing 50 mM HEPES pH 7.2, 2% CTAB (hexadecyltrimethylammonium bromide), 6 M GuHCl (guanidine hydrochloride), and 5 mM DTT. Cells were lysed by sonication: 10 pulses, 30 seconds, at 60% amplitude and further heating the lysate at 60 °C for 20 minutes. Next, 200 μL of lysate from every condition was methanol-chloroform precipitated. Protein concentration was determined using the bicinchoninic acid (BCA) protein assay (Thermo Fisher). Samples were then diluted to 2 M GuHCl with 10 mM EPPS pH 8.5 and digested with 20 ng μL$^{-1}$ LysC (Wako) at room temperature overnight. Samples were further diluted to 0.5 M GuHCl with 10 mM EPPS pH 8.5 and digested with an additional 20 ng μL$^{-1}$ LysC and 10 ng μL$^{-1}$ sequencing-grade trypsin (Promega) at 37 °C for 16 h. The digested samples were dried using a vacuum evaporator at room temperature and taken up in 200 mM EPPS pH 8.0. The multiplexing TMTpro tags[9] were added at a mass ratio of 5:1 tag/peptide to ~ 40 μg of peptide per condition and allowed to react for 2 h at room temperature. The reaction was quenched with 1% hydroxylamine (30 min, RT). Samples from all conditions were combined into one tube, acidified with 5% phosphoric acid (pH<2). The samples are then ultracentrifuged at 100 krcf at 4 °C for an hour to pellet undigested proteins. The supernatants were dried using a vacuum evaporator at room temperature to remove acetonitrile from the labeling step. Dry samples were taken up in HPLC grade water and subjected to medium pH reverse phase prefractionation. Samples are prefractionated with medium pH reverse-phase HPLC (Zorbax 300 Extend C18, 4.6 x 250 mm column, Agilent) with 10 mM ammonium bicarbonate, pH 8.0, using 5% acetonitrile for 17 minutes followed by an acetonitrile gradient from 5% to 30%. Each fraction was dried and resuspended in 100 µL of HPLC water. Fractions were acidified to pH <2 with phosphoric acid and desalted. The samples were resuspended in 1% formic acid to 1 μg $\mu L^{-1}$ and 1 μg of the total combined sample was analyzed with the TMTproC approach.[9]

Samples were analyzed on an EASY-nLC 1200 (Thermo Fisher Scientific) HPLC coupled to an Orbitrap Fusion Lumos mass spectrometer (Thermo Fisher Scientific) with Tune version 3.3. Peptides were separated on an Aurora Series emitter column (25 cm × 75 µm ID, 1.6 µm C18) (Ionopticks, Australia) and held at 60 °C during separation using an in-house built column oven, over 120 min for unfractionated and 90 min for fractionated samples, applying nonlinear acetonitrile gradients at a constant flow rate of 350 nL $min^{-1}$. MS parameter were set as previously described for TMTproC analysis.[9]

The data was analyzed using the Gygi Lab GFY software licensed from Harvard. The detailed description can be found in [10].Thermo Fisher Scientific raw-files were converted to mzXML using ReAdW.exe (http://svn.code.sf.net/p/sashimi/code/). Assignment of MS2 spectra was performed using the SEQUEST[11] algorithm by searching the data against the combined reference proteomes for *Escherichia coli* acquired from Uniprot on 08/2017 along with common contaminants such as human keratins and trypsin. The target-decoy strategy was used to construct a second database of reversed sequences that were used to estimate the false discovery rate on the peptide level.[12] SEQUEST searches were performed using a 20-ppm precursor ion tolerance with the requirement that both N- and C terminal peptide ends are consistent with the protease specificities of LysC and Trypsin. A peptide level MS2 spectral assignment false discovery rate of 1% was obtained by applying the target decoy strategy with linear discriminant analysis.[13] Peptides were assigned to proteins and a second filtering step to obtain a 1% FDR on the protein level was applied.[14] Peptides that matched multiple proteins were assigned to the proteins with the most unique peptides. For all methods, peptides were only considered quantified if the signal to FT noise ratio (S:N) across all channels was greater than 40. Identification of complementary ion peaks, modelling of the isolation window, and deconvolution of the complementary peaks were performed as previously described.[9]

The mass spectrometry proteomics data have been deposited to the ProteomeXchange Consortium via the PRIDE[15] partner repository with the dataset identifier PXD042444.

### Supplemental analysis methods

#### Theoretical monoisotopic decay profiles for protein turnover in chemostat

***Detailed model***

In a chemostat, the volume of liquid inside the vessel stays constant.

$$\frac{d(V)}{dt} = F_0 - DV$$

(1)

At steady state, dV/dt=0, therefore the equation 1 simplifies to

$$F_0 = DV$$

(2)

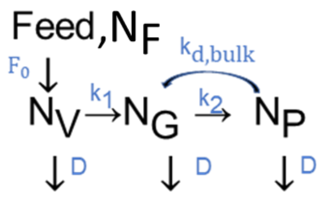


As presented in the schematic above, we assume all nitrogen N in the system is present in three forms: ammonia in the vessel ($N_V$), glutamate that has been assimilated by *E. coli* ($N_G$), and proteins ($N_P$). Ammonia from the feed (concentration $[N_F]$) replenishes ammonia in the vessel ($N_V$) at the volumetric flow rate of $F_0$. The ammonia is assimilated by bacteria in the form of glutamate ($N_G$) with a specific conversion rate of $k_1$. Glutamate is also replenished by the amino acids recycled from degraded proteins at a degradation rate of $k_{d,bulk}$. Glutamate is the precursor for the other amino acids which eventually become proteins ($N_P$) at a rate $k_2$. Each form of nitrogen gets depleted from the vessel at a dilution rate, D.

A balance on total Nitrogen N yields:

$$\frac{dN}{dt} = [N_F]F_0 - DN$$

Where N is the sum of all nitrogen pools. At steady state, dN/dt=0, therefore the above equation simplifies to

$$[N_F]F_0 = DN$$

(3)

A balance on ammonia in the vessel $N_V$ yields:

$$\frac{dN_V}{dt} = [N_F]F_0 - k_1 N_V - DN_V$$

At steady state, $dN_V/dt$, so the above equation simplifies to:

$$[N_F]F_0 = k_1N_V+DN_V \tag{4}$$

Using equation 3 and 4,

$$DN_P+DN_G= k_1N_V$$

A balance on nitrogen in the form of proteins $N_P$ yield:

$$\frac{dN_P}{dt} = k_2N_G - k_{d,bulk}N_P - DN_P$$

At steady state, $dN_P/dt = 0$, so the above equation simplifies to:

$$k_2N_G = k_{d,bulk}N_P + DN_P \tag{5}$$

Let $\mathbf{c = N_P/N_G}$, $\mathbf{d = N_V/N_G}$ **and** $\mathbf{a = (k_{d,bulk} + D)/D}$

Using equations 3, 4, and the above annotations,

$$k_1=D(c+1)/d \tag{6}$$

Using equation 5 and the above annotations,

$$k_2=Dac \tag{7}$$

Next, we determine how the composition of glutamate pools i.e. the amount of light nitrogen in the form of glutamate $N_{GL}$, changes with time. But first we need to determine how the composition of light nitrogen available in the vessel $N_{VL}$ changes with time.

Dynamics of the composition of $N_{VL}$:

$$\frac{d(N_{VL})}{dt} = [N_{FL}]F_0 - k_1N_{VL} - DN_{VL}$$

$[N_{FL}] = 0$ since the feed is switched to heavy nitrogen at t=0. We get,

$$\frac{d(N_{VL})}{dt} = -(k_1+D)\, N_{VL}$$

at t=0, $N_{VL}=N_V$. Integrating the above equation and using the boundary condition,

$$N_{VL}=N_V\, e^{-(k_1+D)\, t} \tag{8}$$

Dynamics of the composition of $N_{GL}$:

$$\frac{d(N_{GL})}{dt} = k_1 N_{VL} + k_{d,bulk} N_{PL} - k_2 N_{GL} - DN_{GL}$$

Using equation 6, 7 and 8,

$$\frac{d(N_{GL})}{dt} = k_1 N_V \, e^{-(k_1+D)\,t} + D(a-1)N_{PL} - D(ac+1)N_{GL} \tag{9}$$

Dynamics of the composition of $N_{PL}$:

$$\frac{d(N_{PL})}{dt} = k_2 N_{GL} - k_{d,bulk} N_{PL} - DN_{PL}$$

Using equation 7

$$\frac{d(N_{PL})}{dt} = DacN_{GL} - DaN_{PL} \tag{10}$$

Equation 9 and 10 form a system of coupled differential equations that can be solved

First solve the homogenous part of the equation. The matrix of interest is

$$\begin{pmatrix} -D(ac+1) & D(a-1) \\ Dac & -Da \end{pmatrix}$$

Eigen values and the corresponding eigen vectors are:

$$\lambda_1 = -D,\ v_1 = \begin{bmatrix} 1 \\ ac/(a-1) \end{bmatrix};\ \lambda_2 = -Da(c+1),\ v_2 = \begin{bmatrix} 1 \\ -1 \end{bmatrix}$$

$$N_{GL} = M_1 e^{-Da(c+1)t} + M_2 e^{-Dt} + M_3 e^{-(k_1+D)\,t} \tag{11}$$

$$N_{PL} = -M_1 e^{-Da(c+1)t} + \frac{M_2 ace^{-Dt}}{a-1}$$

At time 0, $N_{GL} = N_G$ and $N_{PL} = N_P$. We can determine the values of $M_1$ and $M_2$ using the boundary conditions. $M_3$ can be determined using the method of undetermined coefficients.

Finally,

$$M_3 = \frac{N_G\,(c+1)}{dac-c-1};\ M_2 = \left(\frac{N_G\,(c+1)(a-1)}{ac+a-1}\right)\left(\frac{dac-c-2}{dac-c-1}\right);\ M_1 = N_G - M_3 - M_2$$

In a steady state chemostats, total protein amounts (P) per unit cell stays constant. However, the light peak ($M_0$) decays over time as we switch the media from 14N to 15N. For a peptide containing f nitrogen, balance on the mono-isotopic peak ($M_0$):

$$\frac{d(M_0)}{dt} = \text{Rate of synthesis of } M_0 - \text{Rate of removal of } M_0$$

$$\text{Rate of synthesis of } M_0 = \text{Rate of synthesis of total protein} * \text{fraction of light peak}$$

$$= (k_T)\ (\text{Probability of all the nitrogens being light})\ (\text{Probability of all other elements to be light})$$

$$= k_T \left(\frac{N_{GL}}{N_G}\right)^f I$$

$$\text{Rate of removal of } M_0 = \text{First order active decay of } M_0 + \text{Rate of dilution out of the vessel}$$

$$= k_D M_0 + DM_0$$

$$= (k_D + D) M_0$$

Putting together term 1 and term 2:

$$\frac{d(M_0)}{dt} = k_T \left(\frac{N_{GL}}{N_G}\right)^f I - (k_D + D) M_0$$

Since total protein (P) stays constant:

$$\frac{d(P)}{dt} = k_T - (k_D + D) P = 0$$

$$\Rightarrow k_T = (k_D + D) P$$

Additionally at time 0,

$$M_0 = P\ I$$

Using the integrating factor $e^{(k_D+D)t}$

$$\int_0^{M_0(t)} d\left(e^{(k_D+D)t} M_0\right) = \int_0^t k_T I \left(\frac{N_{GL}}{N_G}\right)^f e^{(k_D+D)t} dt$$

$$\frac{M_0}{PI} = e^{-(k_D+D)t} + (k_D + D)\, e^{-(k_D+D)t} \int_0^t \left(\frac{N_{GL}}{N_G}\right)^f e^{(k_D+D)t} dt \tag{12}$$

Using numerical integration, we can get $M_0$ as a function of time.

Now we need to determine estimates for the constants a, c, and d. For P-lim and C-lim cells, the cells glutamate levels are ~ 100 mM and protein levels are ~1000 mM.[16]

$$c = N_P/N_G = 10\,.$$

The theoretical decay profiles of $M_0$ are only sensitive to lower values of c.

The concentration of nitrogen in the feed for P-lim and C-lim cells is 9.5 mM and for N-lim cells is 1.9 mM. Since under N-lim, the cells consume all the nitrogen provided and cells grow to approximately the same $OD_{600}$ in each condition, we assume that 1.9 mM out of the 9.5 mM is consumed by bacteria under P-lim and C-lim.

$$d = N_V/N_G = \frac{(9.5-1.9)*(c+1)}{1.9} = \frac{(9.5-1.9)*(11)}{1.9} = 44$$

Since there is minimal degradation under P-lim, we assume that the recycling flux is 0

$$a = (k_{d,bulk} + D)/D = 1$$

Under N-lim, glutamate pools must be very small, therefore, c is set to be 100 (arbitrarily large).

Since the pools of nitrogen outside the vessel are 0, $N_V = 0 \Rightarrow M_3 = \frac{N_G\,(c+1)}{dac\text{-}c\text{-}1} = \frac{k_1 N_V}{Dac\text{-}k_1} = 0$

Assuming a recycling rate of 0.05 $hr^{-1}$(calculated from Fig. 3G), a= 1.38 for 6 hours doubling chemostat.

***Simplified model***

Based on our estimate of the constants a,c,d; we deduced that the above model can be greatly simplified in its mathematical form.

For instance, equation 9 can be simplified as follows. The dynamics for the composition of nitrogen available for the cells to make proteins can be written as:

$$\frac{N_L}{N_T} = e^{-DKt}$$

Where $K \in [1, \infty\}$ is a constant, which is set large (K=100) for N-lim cells, indicating that the exchange of light to heavy nitrogen is quite fast in N-lim cells. However, K is set to be 1 for P-lim and C-lim cells, indicating that the major portion of nitrogen is present outside the cells and is only exchanged by dilution out of the vessel and not by bacterial consumption. Balance on the mono-isotopic peak ($M_0$) for the simplified model:

$$\frac{d(M_0)}{dt} = \text{Rate of synthesis of } M_0 - \text{Rate of removal of } M_0$$

Rate of synthesis of $M_0$= Rate of synthesis of total protein * fraction of light peak

= ($k_T$) (Probability of all the nitrogens being light) (Probability of all other elements to be light)

$$= k_T \left(\frac{N_L}{N_T}\right)^f I$$

Rate of removal of $M_0$= First order active decay of $M_0$ + Rate of dilution out of the vessel

$$= k_D M_0 + DM_0$$

$$= (k_D+D)M_0$$

Putting together term 1 and term 2:

$$\frac{d(M_0)}{dt} = k_T \left(\frac{N_L}{N_T}\right)^f I - (k_D+D)M_0$$

Since total protein (P) stays constant:

$$\frac{d(P)}{dt} = k_T - (k_D+D)P = 0$$

$$=> k_T = (k_D+D)P$$

Additionally at time 0,

$$M_0 = P\ I$$

Substituting ths simplified expression of $\frac{N_L}{N_T}$ from above

$$\frac{d(M_0)}{dt} = k_T \left(\frac{N_L}{N_T}\right)^f I - (k_D+D)M_0$$

$$= (k_D+D)\ (PI\ e^{-DKf\ t} - M_0)$$

Integrating using the integrating factor $e^{(K_D+D)t}$ and applying the boundary condition; we obtain an analytical solution for the fraction of $M_0$ over time.

$$\frac{M_0}{P} = \frac{(k_D+D)\ e^{-fDKt} - fDK\ e^{-(k_D+D)\ t}}{(k_D+D) - fDK} \qquad (13)$$

Equation 12 and Equation 13 represent the detailed and simplified form of the model respectively. Figure S3 shows the equivalence for the two models in all the conditions and hence justifies the use of the simplified model. Figure S3A plots N-limited cells (parameters c=100, a=1.38, $M_3 = 0$, nitrogen=15), and S3B plots P-limited cells (parameters c=10, a=1.07, d= 44, nitrogen=15).

#### Protein half-life fitting

Due to the pipetting errors, the observed data needs to be normalized. We assume that the mode of the membrane peptides (annotations from Uniprot) is stable and use it for normalization. To remove membrane peptides which are actively degrading, we applied the K-means clustering algorithm to cluster the peptides according to their $M_0$ decay profiles over time. All the membrane peptides, except the ones that clustered into the most rapidly degrading profiles, were used for normalization. The peptides were divided into g groups based on the number of nitrogen atoms (f) the constituent peptides contain. To avoid forcing the time 0 data point to the value of 1, we used the fractional form of the theoretical M0 decay (Eq. 13) by dividing the value at each time point with the sum of values from the 8 time points. Accordingly, the fractional form of experimental data was also obtained.

The correction values for each time point were obtained by minimizing the least square difference between the theoretical value (equation 13 with $K_D$=0) and the median values of the experimentally observed $M_0$ for all the g groups and 8 time points at once. Gradient descent (curve_fit function in python) was used to find the minima.

- Fractional form of the theoretical $M_0$ ($\widetilde{M0}_{theoretical,\ g,\ t}$) is given by $\frac{M0_{theoretical,g,t}}{\sum_t M0_{theoretical,g,t}}$
- Median of the fractional form of the experimental $M_0$ (mdn $\widetilde{M0}_{experimental,\ g,\ t}$) is given by $\underset{k}{\text{median}} \frac{M0_{experimental,g,t,k}}{\sum_t M0_{experimental,g,t,k}}$, where k corresponds to all the peptides in a group g.
- a is a vector of length 8 and $a_t$ denotes an element in the vector.

$$\hat{a} \triangleq \arg\min_a \sum_g \sum_{t=0}^{7} (\widetilde{M0}_{theoretical,\ g,\ t}(f) - a_t\ (\text{mdn}\ \widetilde{M0}_{experimental,\ g,\ t}))^2$$

Once normalized, the data from different peptides of a protein is fit to obtain a protein-specific $k_D$. To avoid forcing the time 0 data point to the value of 1, we used the fractional form of the theoretical equation by dividing the value at each time point with the sum of values from the 8 time points. For a protein p with i peptides each containing f number of nitrogens, we construct a theoretical matrix of (px8) values. Accordingly, the corresponding matrix of experimental values is created. The $k_D$ for each time point is obtained by minimizing the least square difference between the theoretical value (equation 13 with a free parameter $k_D$) and the experimentally observed values. Gradient descent (curve_fit function in python) was used to find the minima.

- Fractional form of the theoretical $M_0$: $\widetilde{M0}_{theoretical,\ p,\ i,\ t}(k_D,f)$ is given by $\frac{M0_{theoretical,\ p,\ i,\ t}(k_D,f)}{\sum_t M0_{theoretical,\ p,\ i,\ t}(k_D,f)}$
- Fractional form of the experimental $M_0$: $\widetilde{M0}_{experimental,\ p,\ i,\ t}$ is given by $\frac{M0_{experimental,\ p,\ i,\ t}}{\sum_t M0_{experimental,\ p,\ i,\ t}}$

$$\widehat{k_{D,p}} \triangleq \arg\min_{k_{D,p}} \sum_i \sum_{t=0}^{7} (\widetilde{M0}_{theoretical,\ p,\ i,\ t}(k_D,f) - \widetilde{M0}_{experimental,\ p,\ i,\ t})^2$$

To assign confidence to actively degrading proteins. For bacteria doubling with 6 hours $H_0$: $T_{1/2} = 6$, $H_1$: $T_{1/2} < 6$ and the test statistic $t = (\bar{x}-6)/\sqrt{\frac{\sigma^2}{2}}$ where $\bar{x}$ is the sample mean and $\sigma^2$ is the sample variance calculated using the two replicates. *p*-values are assigned using one-tailed t-test. We called the proteins with *p*-value < 0.05 as actively degrading.

#### Cumulative fold enrichment calculation

For the molecular weight case the cumulative log enrichment for a particular half-life is defined as:

$$\mathrm{Log2}\left(\frac{\text{\# of small proteins (Mwt < 10 KDa) < half-life}}{\text{\# of large proteins(Mwt} \geq \text{10 KDa)< half-life}} \times \frac{\text{Total \# of large proteins (Mwt} \geq \text{10 KDa)}}{\text{Total \# of small proteins (Mwt < 10 KDa)}}\right)$$

Similarly, for the intrinsically disordered case the cumulative log enrichment for a particular half-life is defined as:

$$\mathrm{Log2}\left(\frac{\text{\# of disordered proteins (}\geq\text{ 50\% disorder) < half-life}}{\text{\# of ordered proteins (< 50\% disorder)< half-life}} \times \frac{\text{Total \# of ordered proteins (< 50\% disorder)}}{\text{Total \# of disordered proteins (}\geq\text{ 50\% disorder)}}\right)$$

#### Likelihood ratio calculations for the constant and the scaled model

Both the constant and the scaled model describe the relationship between $T_{1/2}$ total for bacteria doubling with 12 hours and a shorter doubling time of x h. Assuming that the errors around both the scaled model and the non-scaled model are normally distributed with a fixed noise (σ), the likelihood can be given as:

Scaled Model: $T_{1/2,\,x} = rT_{1/2,\,12} + \varepsilon_{scaled}$ , r is a constant

$$p\left(\varepsilon_{scaled} \,\middle|\, T_{1/2,\,x}, T_{1/2,\,12}, \text{scaled model}, \sigma_{scaled}\right) = N(0|\mu=(T_{1/2,\,x} - rT_{1/2,\,12}), \sigma_{scaled}^2)$$

Constant Model: $T_{1/2,\,x} = g(T_{1/2,\,12}) + \varepsilon_{constant}$

$$p\left(\varepsilon_{constant} \,\middle|\, T_{1/2,x}, T_{1/2,\,12}, \text{constant model}, \sigma_{constant}\right) = N\,(0|\mu=(T_{1/2,\,x} - g(T_{1/2,\,12})), \sigma_{constant}^2)$$

Estimating the parameter σ by maximizing the probability functions with respect to it:

If n is the total number of proteins, likelihood (L) is given by:

$$L = \prod_{i=1}^{n} p(0|\,\mu, \sigma^2)$$

$$\text{Log}(L) = \frac{-n}{2}\log(2\pi) - n\log(\sigma) - \frac{1}{2\sigma^2}\sum_{i=1}^{n}\mu^2$$

Taking the derivative wrt to σ and equating to zero:

$$\hat{\sigma}^2 = \frac{\sum_{i=1}^{n}\mu^2}{n}$$

and substituting it the ratio of the likelihoods for the scaled and the constant model as follows:

$$\frac{\hat{L}_{CONSTANT}}{\hat{L}_{SCALED}} = \exp(-n/2(\ln(RSS_{constant}) - \ln(RSS_{scaled})))$$

$$RSS_{constant} = \sum_i (T_{1/2,\,x,\,i} - rT_{1/2,\,12,i})^2$$

$$RSS_{scaled} = \sum_i (T_{1/2,\,x,\,i} - g(T_{1/2,\,12,i}))^2$$

where the index i is for the proteins and n is the total number of proteins.

#### Assigning substrates to proteases

To identify the proteins that get stabilized on knocking out a protease, we compared the half-lives ($T_{1/2}$) in the wild-type (WT) and the knockout (KO) strain. $H_0$: $T_{1/2,\,WT} = T_{1/2,\,KO}$, $H_1$: $T_{1/2,\,WT} < T_{1/2,\,KO}$ and the test statistic $t = (\bar{x}_{WT} - \bar{x}_{KO}) / \sqrt{\frac{\sigma_{WT}^2 + \sigma_{KO}^2}{2}}$ where $\bar{x}$ is the sample mean and $\sigma^2$ is the sample variance calculated using replicates. *p*-values are assigned using one-tailed t-tests. We called the proteins with *p*-value < 0.10 as confidently stabilized.

For the substrates that are confidently stabilized in the triple KO, we could assign the percentage of contribution towards stability by each of the 6 mutually exclusive categories: ClpP alone, Lon alone, HslV alone, additive contribution, redundant contribution and actively degrading in triple knockout. All the calculations are done on the rates ($k = \ln(2)/T_{1/2}$) and not on half-lives.

Gain in stability by each individual knockout is defined as

$$\Delta k_{KO} = \begin{cases} k_{WT} - k_{KO} & \text{if } k_{WT} > k_{KO} \\ 0 & \text{otherwise} \end{cases}$$

If D is the bacterial growth rate, percentage contribution of individual knockouts towards complete stability is defined as:

$$\% \text{ KO} = \frac{\Delta k_{KO}}{k_{WT} - D}$$

$$\% \text{ Additive} = \frac{\Delta k_{\Delta clpP} + \Delta k_{\Delta lon} + \Delta k_{\Delta hslV}}{k_{WT} - D}$$

$$\% \text{ Redundant} = \frac{\Delta k_{\Delta clpP \Delta lon \Delta hslV} - (\Delta k_{\Delta clpP} + \Delta k_{\Delta lon} + \Delta k_{\Delta hslV})}{k_{WT} - D}$$

$$\% \text{ Unexplained} = \frac{(k_{WT} - D) - \Delta k_{\Delta clpP \Delta lon \Delta hslV}}{k_{WT} - D}$$

These percentages are further modified so that any negative values are set to 0. The modified percentages are renormalized by dividing them with the sum of all the categories.

We assigned each confidently degrading protein to one of six categories: clpP substrate, lon substrate, hslV substrate, additively degraded by multiple proteases, redundantly degraded by multiple proteases, and unexplained. We assigned each protein using the following criteria in order (Redistribute the proteins again in unexplained vs the redundant category):

1. Proteins were assigned as a substrate of a single protease if at least 60% of the increase in stability is explained by the corresponding knock-out.

2. If the individual contributions of each protease summed to at least 70%, the protein was categorized as additive.

3. If more than 20% was explained by the redundancy, the proteins were categorized as redundant.

4. All the remaining proteins and the ones that are still short-lived in the Δ*clpP*Δ*lon*Δ*hslV* were categorized as unexplained.

#### Percentage of active proteome turnover per unit hour

Absolute protein abundances ($P_a$) are calculated using label free mass spectrometry (Table S6). The proteins with half-lives greater than the doubling time are called stable and assigned a half-life equal to the doubling time ($T_{1/2,\ cap}$). Percentage of active proteome turnover is calculated by $\frac{\sum_i MW_i \frac{\ln(2)}{T_{1/2,cap,i}} P_{a,i} - \sum_i MW_i\, D\, P_{a,i}}{\sum_i MW_i\, P_{a,i}}$ where i is an index for proteins and D is the bacterial growth rate.
